# Ice2 promotes ER membrane biogenesis in yeast by inhibiting the conserved lipin phosphatase complex

**DOI:** 10.1101/2020.02.23.961722

**Authors:** Dimitrios Papagiannidis, Peter W. Bircham, Christian Lüchtenborg, Giulia Ruffini, Britta Brügger, Sebastian Schuck

## Abstract

Cells dynamically adapt organelle size to current physiological demand. Organelle growth requires membrane biogenesis and therefore needs to be coordinated with lipid metabolism. The endoplasmic reticulum (ER) can undergo massive expansion, but the regulatory mechanisms that govern ER membrane biogenesis are largely unclear. Here, we conduct a genetic screen for factors involved in ER membrane expansion in budding yeast and identify the ER transmembrane protein Ice2 as a strong hit. We show that Ice2 promotes ER membrane biogenesis by opposing the phosphatidic acid phosphatase Pah1, which is called lipin in metazoa. Ice2 inhibits the conserved Nem1-Spo7 complex, which dephosphorylates and thus activates Pah1. Furthermore, Ice2 cooperates with the transcriptional regulation of lipid synthesis genes and helps to maintain ER homeostasis during ER stress. These findings establish inhibition of the lipin phosphatase complex as an important mechanism for controlling ER membrane biogenesis.

## INTRODUCTION

Cells reshape and resize their organelles when they undergo differentiation or adapt to changing environmental conditions. An increase in organelle size typically involves enhanced membrane biogenesis, which in turn requires an adequate supply of membrane lipids. As a result, organelle biogenesis depends on whether cells employ available lipids as building blocks for new membranes, consume them for energy production or store them for future use. Accordingly, the regulatory mechanisms that control lipid synthesis and utilization are fundamental for organelle biogenesis.

The ER is a morphologically complex organelle with essential functions in protein folding and lipid synthesis (Westrate et al., 2015). It forms the nuclear envelope and extends into the cytoplasm as a network called the peripheral ER. The principal structural elements of the ER are tubules and sheets (Shibata et al., 2010). In addition, intermediate structures exist, such as tubular matrices and sheets with fenestrations (Puhka et al., 2012; Nixon-Abell et al., 2016; Schroeder et al., 2019). A variety of organelle morphologies can arise according to physiological demand, ranging from mainly tubular ER in lipid hormone-producing cells of the testes to mainly sheet-like ER in secretory cells of the pancreas (Fawcett, 1981). Besides morphology, ER size is also tuned to cellular need. For instance, the ER expands several-fold when B lymphocytes differentiate into antibody-secreting plasma cells or when cells face protein folding stress in the ER (Wiest et al., 1990; Bernales et al., 2006). Such stress-induced ER expansion is mediated by the unfolded protein response (UPR), which induces genes encoding ER-resident protein folding enzymes to restore homeostasis (Walter and Ron, 2011). In addition to determining the abundance of protein folding enzymes, the UPR also regulates ER membrane biogenesis. It does so, at least in part, by inducing genes that encode lipid synthesis enzymes (Sriburi et al., 2007; Bommiasamy et al., 2009; Schuck et al., 2009).

Yeast synthesize membrane phospholipids primarily from phosphatidic acid (PA) through the CDP-DAG pathway (Henry et al., 2012). Many enzymes of this pathway are controlled transcriptionally by the activators Ino2/4 and the repressor Opi1. Ino2 and Ino4 form a heterodimer that binds to promoter elements of lipid synthesis genes. Opi1 inhibits Ino2/4 by binding to Ino2 (Heyken et al., 2005). Repression of Ino2/4 by Opi1 is relieved when accumulating PA tethers Opi1 to the ER membrane, sequestering it away from the nucleoplasm (Loewen et al., 2004). Thus, the PA-Opi1- Ino2/4 system forms a feedback loop and matches PA availability to the cellular capacity for converting PA into other phospholipids. Removal of Opi1 results in activation of lipid synthesis and ER membrane expansion, even in cells lacking the UPR. This membrane expansion without a corresponding upregulation of the protein folding machinery increases cellular resistance to ER stress, highlighting the physiological importance of ER membrane biogenesis (Schuck et al., 2009). However, it is unknown whether activation of Ino2/4 is the only mechanism regulating the production of ER membrane. Furthermore, Ino2/4 and Opi1 are not conserved in metazoa. Therefore, lipid metabolism could be regulated in unique ways in yeast. Alternatively, conserved regulators of lipid metabolism distinct from Ino2/4 and Opi1 could determine ER size in both yeast and higher eukaryotes.

Here, we systematically search for genes involved in ER membrane biogenesis in budding yeast, *Saccharomyces cerevisiae*, to better define the regulatory circuitry that connects lipid metabolism and organelle biogenesis.

## RESULTS

### An inducible system for ER membrane biogenesis

Removal of Opi1 induces Ino2/4-driven lipid synthesis genes and thereby leads to expansion of the ER (Schuck et al., 2009). To improve experimental control over ER membrane biogenesis, we developed an inducible system using ino2(L119A), an Ino2 variant that cannot be inhibited by Opi1 (Heyken et al., 2005). We placed ino2(L119A), here termed ino2*, under the control of the *GAL* promoter and employed an expression system that activates this promoter upon addition of the metabolically inert sterol ß- estradiol (Pincus et al., 2014). High-level expression of ino2* is expected to displace endogenous Ino2 from the promoters of its target genes, stimulate lipid synthesis and drive ER membrane biogenesis (Figure 1A; Schuck et al., 2009). Fluorescence microscopy confirmed that expression of ino2* triggered pronounced ER expansion (Figure 1B). In untreated cells, the peripheral ER mostly consisted of tubules, which appeared as short membrane profiles along the cell cortex in optical mid sections and as a network in cortical sections. In contrast, estradiol-treated cells had a peripheral ER that predominantly consisted of ER sheets, as evident from long membrane profiles in mid sections and solid membrane areas in cortical sections.

**Figure 1.**
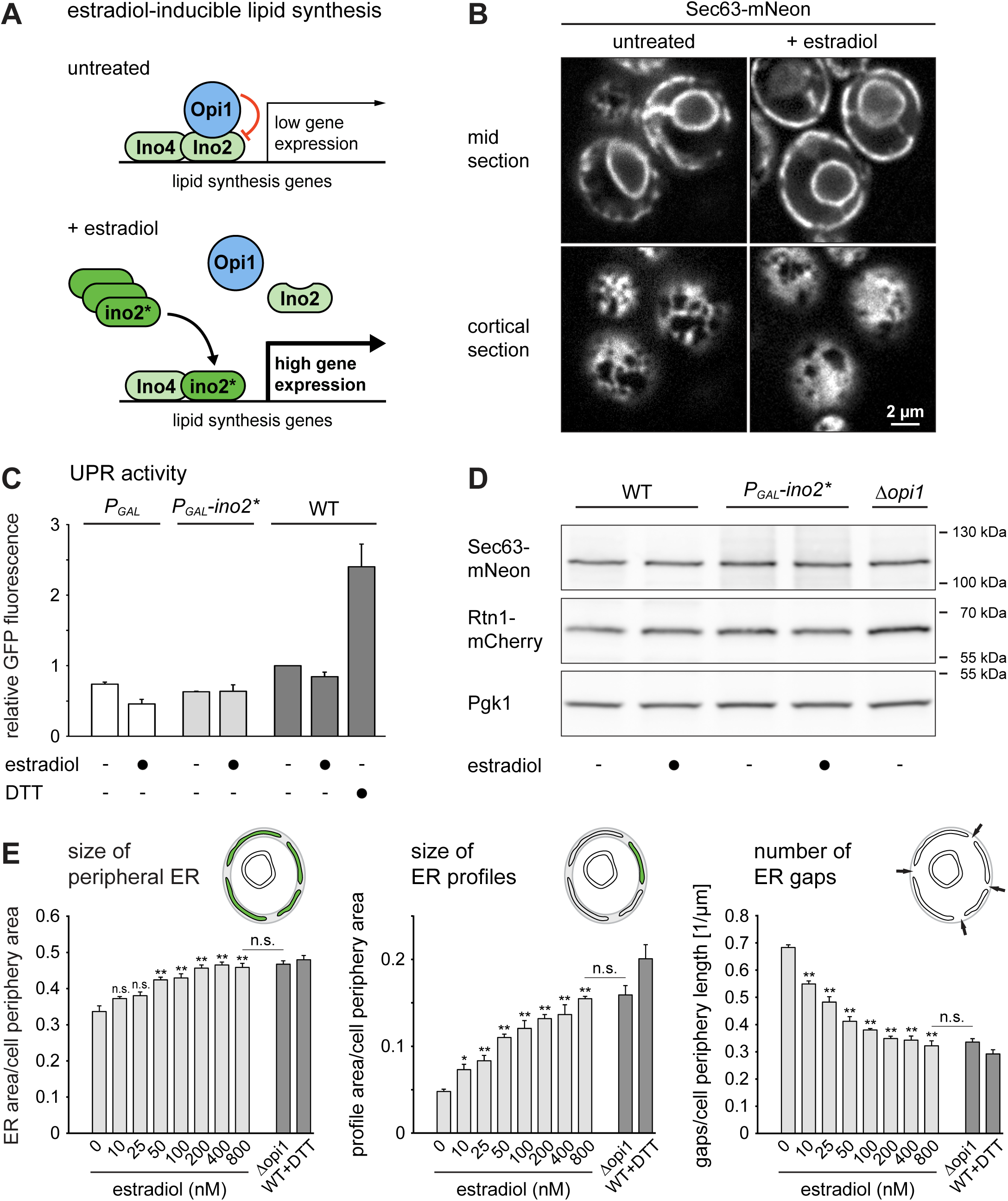
An inducible system for ER membrane biogenesis. **(A)** Schematic of the control of lipid synthesis by estradiol-inducible expression of ino2*. **(B)** Sec63-mNeon images of mid and cortical sections of cells harboring the estradiol-inducible system (SSY1405). Cells were untreated or treated with 800 nM estradiol for 6 h. **(C)** Flow cytometric measurements of GFP levels in cells containing the UPR reporter. WT cells containing the UPR reporter (SSY2306), cells additionally harboring an estradiol- inducible *GAL* promoter (SSY2307) and cells additionally harboring the system for estradiol-inducible expression of ino2* under the *GAL* promoter (SSY2308) were untreated, treated with 800 nM estradiol for 6 h or treated with 8 mM DTT for 1 h. Data were normalized to untreated WT cells. Bars represent the mean of three independent experiments (n = 3), error bars are the standard error of the mean (SEM). WT, wild- type. **(D)** Western blot of Sec63, mCherry and Pgk1 from WT cells (SSY1404), cells harboring the estradiol-inducible system (SSY1405) or *Δopi1* cells (SSY1607), all of which expressed Sec63-mNeon and Rtn1-mCherry. Cells were untreated or treated with 800 nM estradiol for 6 h. Pgk1 served as a loading control. **(E)** Quantification of ER size in estradiol-treated cells harboring the inducible system (SSY1405), untreated *Δopi1* cells (SSY1607) and WT cells (SSY1404) treated with 8 mM DTT for 1 h. Bars represent mean ± SEM, n = 3. Asterisks indicate statistical significance compared with 0 nM estradiol or *Δopi1* cells as determined with a two-tailed Student’s t-test. *, p < 0.05; **, p < 0.01; n.s., not significant.

To test whether ino2*-induced ER expansion causes ER stress, we measured UPR activity by means of a transcriptional reporter. This reporter is based on UPR response elements controlling expression of GFP (Jonikas et al., 2009). Cell treatment with the ER stressor DTT activated the UPR reporter, as expected, whereas expression of ino2* from the *GAL* promoter did not (Figure 1C). Furthermore, expression of ino2* or removal of Opi1 did not alter the abundance of the chromosomally tagged ER proteins Sec63-mNeon or Rtn1-mCherry, even though the *SEC63* gene is regulated by the UPR (Figure 1D; Pincus et al., 2014). These observations indicate that ino2*-induced ER expansion is not an indirect consequence of ER stress but directly arises from enhanced lipid synthesis.

To assess ER membrane biogenesis quantitatively, we developed three metrics for the size of the peripheral ER at the cell cortex, as visualized in mid sections: (1) total size of the peripheral ER, (2) size of individual ER profiles and (3) number of gaps between ER profiles (Figure 1E). These new metrics are less sensitive to variation in image quality than the index of expansion we had used previously (Schuck et al., 2009). Measurements after induction of ino2* expression with different concentrations of estradiol showed a dose-dependent increase in peripheral ER size and ER profile size, and a decrease in the number of ER gaps. After treatment with 800 nM estradiol, the ER was indistinguishable from that in *Δopi1* cells, and we used this concentration in subsequent experiments.

These results show that the inducible system allows titratable control of ER membrane biogenesis without causing ER stress.

### A genetic screen for regulators of ER membrane biogenesis

To identify genes involved in ER expansion, we introduced the inducible ER biogenesis system and the ER markers Sec63-mNeon and Rtn1-mCherry into a knockout strain collection. This collection contains single gene deletion mutants for most of the approximately 4800 non-essential genes in yeast (Giaever et al., 2002). We induced ER expansion by ino2* expression and acquired images by automated microscopy. Based on inspection of Sec63-mNeon in optical mid sections, we defined six phenotypic classes. Mutants were grouped according to whether their ER was (1) underexpanded, (2) properly expanded and morphologically normal, (3) overexpanded, (4) overexpanded with extended cytosolic sheets, (5) overexpanded with disorganized cytosolic structures, or (6) clustered. Figure 2A shows two examples of each class. To refine the search for mutants with an underexpanded ER, we applied the three ER size metrics described above (Figure 2B and Table S1A). This computational analysis confirmed the underexpansion mutants identified visually and retrieved a number of additional, weaker hits.

**Figure 2.**
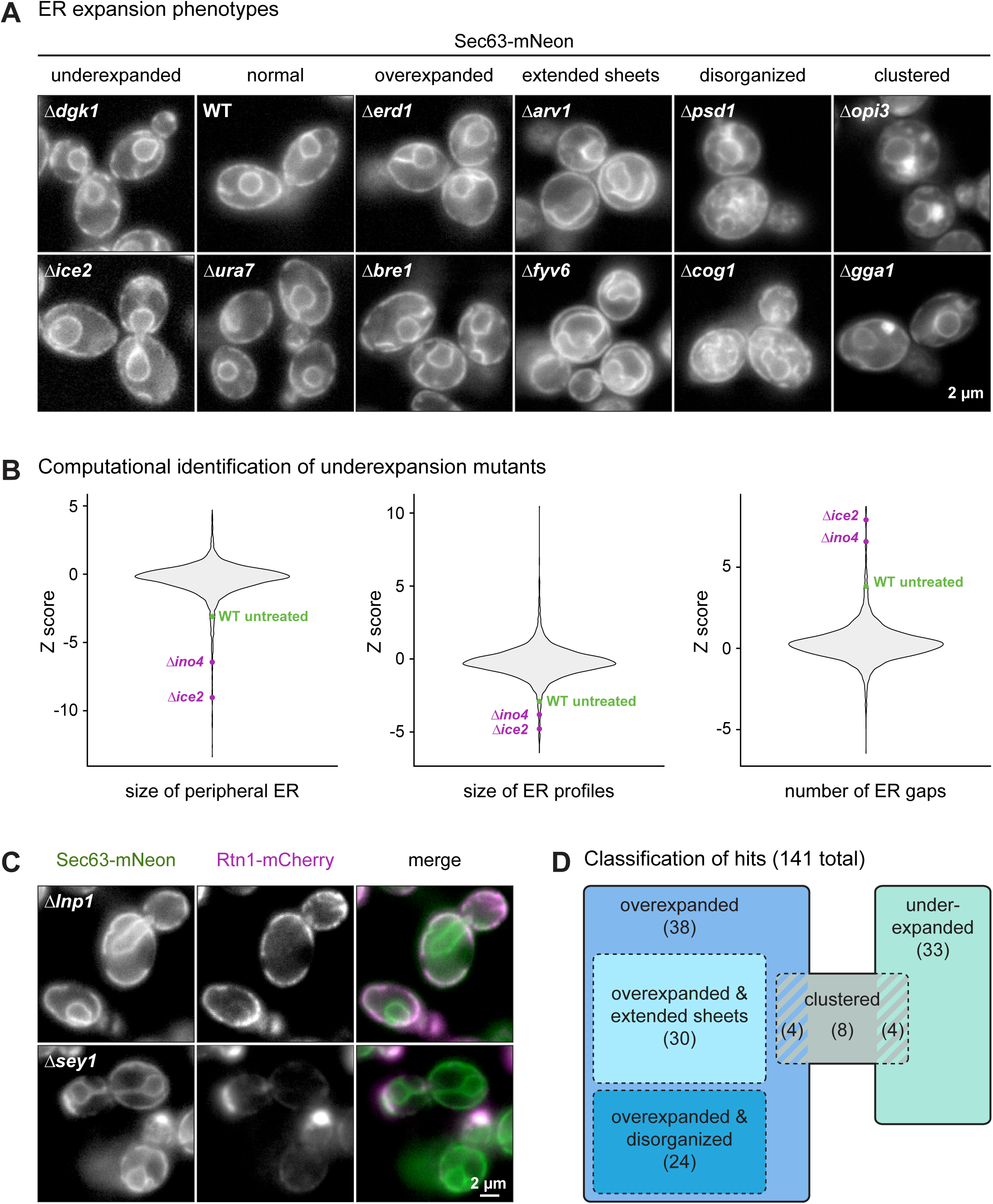
A genetic screen for factors involved in ER membrane biogenesis. **(A)** Sec63-mNeon images of cells of the indicated genotypes harboring the inducible system. Cells were treated with 800 nM estradiol for 6 h. Two examples of each phenotypic class are shown. **(B)** Violin plots of Z scores from the three metrics for ER size determined for each of the 4800 mutant strains. The untreated WT is shown for reference. **(C)** Sec63-mNeon and Rtn1-mCherry images of *Δlnp1* and *Δsey1* cells harboring the inducible system and treated with 800 nM estradiol for 6 h. **(D)** Classification of hits. Numbers in brackets indicate the number of mutants in each class. Striped areas indicate mutants belonging to two classes.

In total, we found 141 mutants that fell into at least one phenotypic class other than morphologically normal (Table S1B). Hits included mutants lacking the ER-shaping gene *LNP1*, which had an overexpanded cortical ER with large gaps, and mutants lacking the *SEY1* gene important for homotypic ER fusion, which displayed ER clusters (Figure 2C; Hu et al., 2009; Chen et al., 2012). The re-identification of these known ER morphogenesis genes validated our approach. About two thirds of the identified mutants had an overexpanded ER, one third had an underexpanded ER and some mutants showed ER clusters together with an otherwise underexpanded, normal or overexpanded ER (Figure 2D). Overexpansion mutants were enriched in gene deletions that activate the UPR (Table S1C; Jonikas et al., 2009). This enrichment suggested that ER expansion in these mutants resulted from ER stress rather than enforced lipid synthesis. Indeed, re-imaging of the overexpansion mutants revealed that their ER was expanded already without ino2* expression (data not shown). Underexpansion mutants included those lacking *INO4* or the lipid synthesis genes *OPI3*, *CHO2* and *DGK1*. In addition, mutants lacking *ICE2* showed a strong underexpansion phenotype (Figure 2A, B).

Overall, our screen indicates that a large number of genes impinge on ER membrane biogenesis, as might be expected for a complex biological process. The functions of many of these genes in ER biogenesis remain to be uncovered. Here, we follow up on *ICE2* because of its critical role in building an expanded ER. Ice2 is a polytopic ER membrane protein (Estrada de Martin et al., 2005) but does not possess obvious domains or sequence motifs that provide clues to its molecular function.

### Ice2 promotes ER membrane biogenesis

To more precisely define the contribution of Ice2 to ER membrane biogenesis, we analyzed optical sections of the cell cortex. Well-focused cortical sections are more difficult to acquire than mid sections but provide more morphological information. Qualitatively, deletion of *ICE2* had little effect on ER structure at steady state but severely impaired ER expansion upon ino2* expression (Figure 3A).

**Figure 3.**
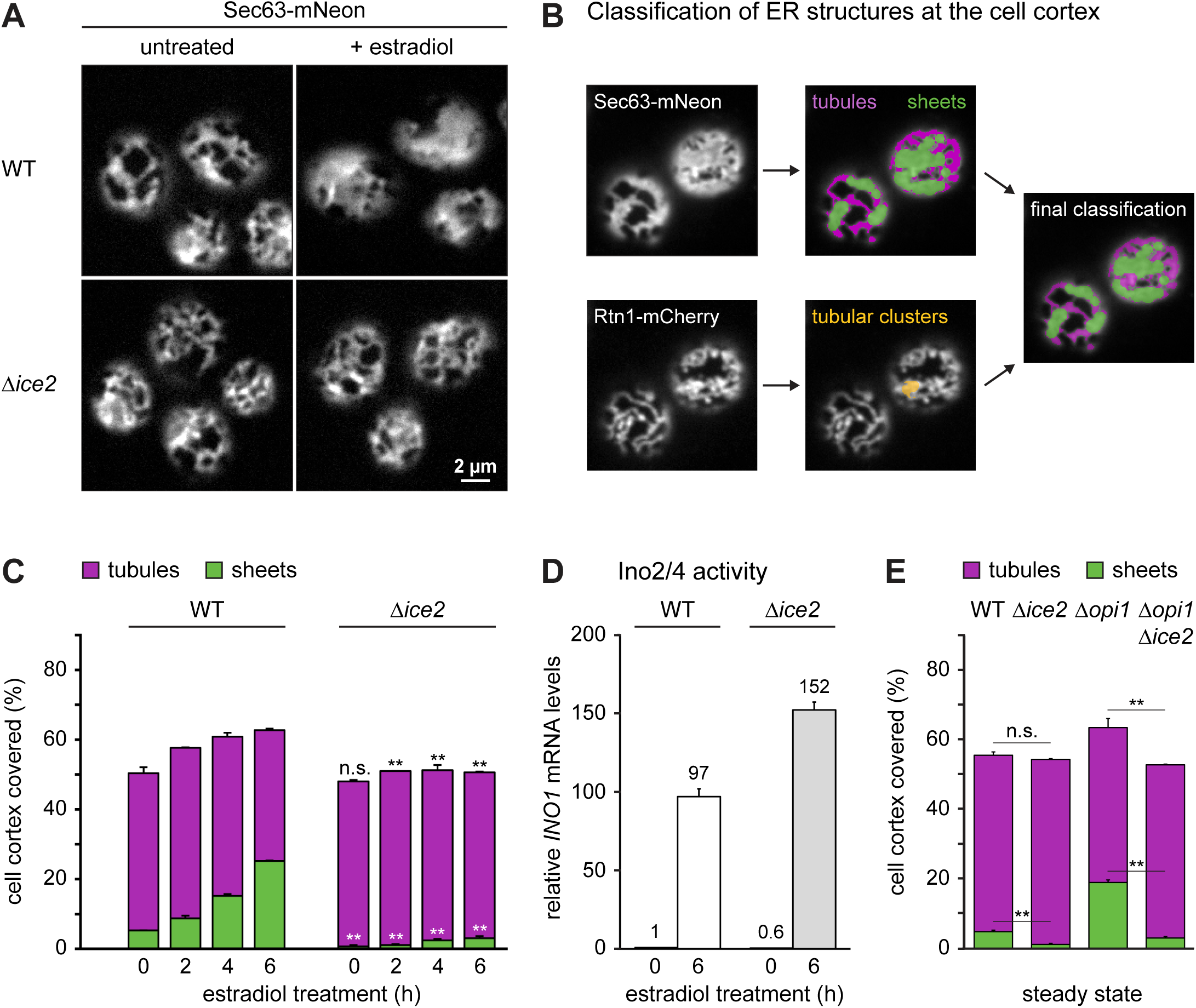
Ice2 is required for ER membrane biogenesis upon activation of Ino2/4. **(A)** Sec63-mNeon images of the cortical ER of WT and *Δice2* cells harboring the inducible system (SSY1405, 1603). Cells were untreated or treated with 800 nM estradiol for 6 h. **(B)** Classification of peripheral ER structures from cortical sections of cells expressing Sec63-mNeon and Rtn1-mCherry as tubules (purple), sheets (green) or tubular clusters (yellow). Tubular clusters are combined with tubules in the final classification, as illustrated by the overlay. **(C)** Quantification of peripheral ER structures in WT and *Δice2* cells harboring the inducible system (SSY1405, 1603) and treated with 800 nM estradiol for the times indicated. Plotted is the mean percentage of cell cortex covered by tubules (purple) or sheets (green), n = 3. Upper error bars are SEM for the sum of tubules and sheets, lower error bars are SEM for sheets. Asterisks indicate statistical significance compared with the corresponding value in WT cells. **, p < 0.01; n.s., not significant. **(D)** mRNA levels of the Ino2/4 target gene *INO1* upon ino2* expression in WT and *Δice2* cells harboring the inducible system (SSY1405, 1603) as measured by quantitative real-time PCR. Data were normalized to untreated WT cells. Mean ± SEM, n = 3. **(E)** Quantification of peripheral ER structures in untreated WT, *Δice2*, *Δopi1* and *Δice2 Δopi1* cells (SSY1404, 2356, 2595, 2811). Bars, error bars and asterisks are as in panel C.

To describe ER morphology quantitatively, we developed a semi-automated algorithm that classifies ER structures as tubules or sheets based on images of Sec63-mNeon and Rtn1-mCherry in cortical sections (Figure 3B). First, the image of the general ER marker Sec63-mNeon is used to segment the entire ER. Second, morphological opening, that is the operation of erosion followed by dilation, is applied to the segmented image to remove narrow structures. The structures removed by this step are defined as tubules and the remaining structures are provisionally classified as sheets. Third, the same procedure is applied to the image of Rtn1-mCherry, which marks high-curvature ER (Westrate et al., 2015). Rtn1 structures that remain after morphological opening and overlap with persistent Sec63 structures are termed tubular clusters. These structures appear as sheets in the Sec63 image but the overlap with Rtn1 identifies them as tubules. Tubular clusters made up only a minor fraction of the total ER and may correspond to so-called tubular matrices observed in mammalian cells (Nixon-Abell et al., 2016). Last, for a simple two-way classification, tubular clusters are added to the tubules and any remaining Sec63 structures are defined as sheets. This analysis using a general and a high-curvature ER marker allows to distinguish densely packed tubules from sheets.

The algorithm described above showed that the ER covered approximately 50% of the cell cortex in untreated wild-type cells and consisted mostly of tubules, as reported (Figure 3C; Schuck et al., 2009; West et al., 2011). Expression of ino2* triggered ER expansion by stimulating the formation of sheets. *Δice2* cells had a defect in sheet formation already at steady state and membrane expansion upon ino2* expression failed almost completely. Importantly, activation of the prototypic Ino2/4 target gene *INO1* upon ino2* expression was intact in *Δice2* cells, ruling out that *ICE2* deletion disrupted the inducible ER biogenesis system (Figure 3D). In addition, *ICE2* deletion abolished the constitutive ER expansion in *Δopi1* cells, excluding that the expansion defect in *Δice2* cells merely reflected a delay (Figure 3E).

Next, we tested whether Ice2 was required for ER expansion induced by ER stress. DTT treatment of wild-type cells triggered rapid ER expansion, which was again driven by the formation of sheets (Figure 4A). Expansion was retarded in *Δice2* cells.

**Figure 4.**
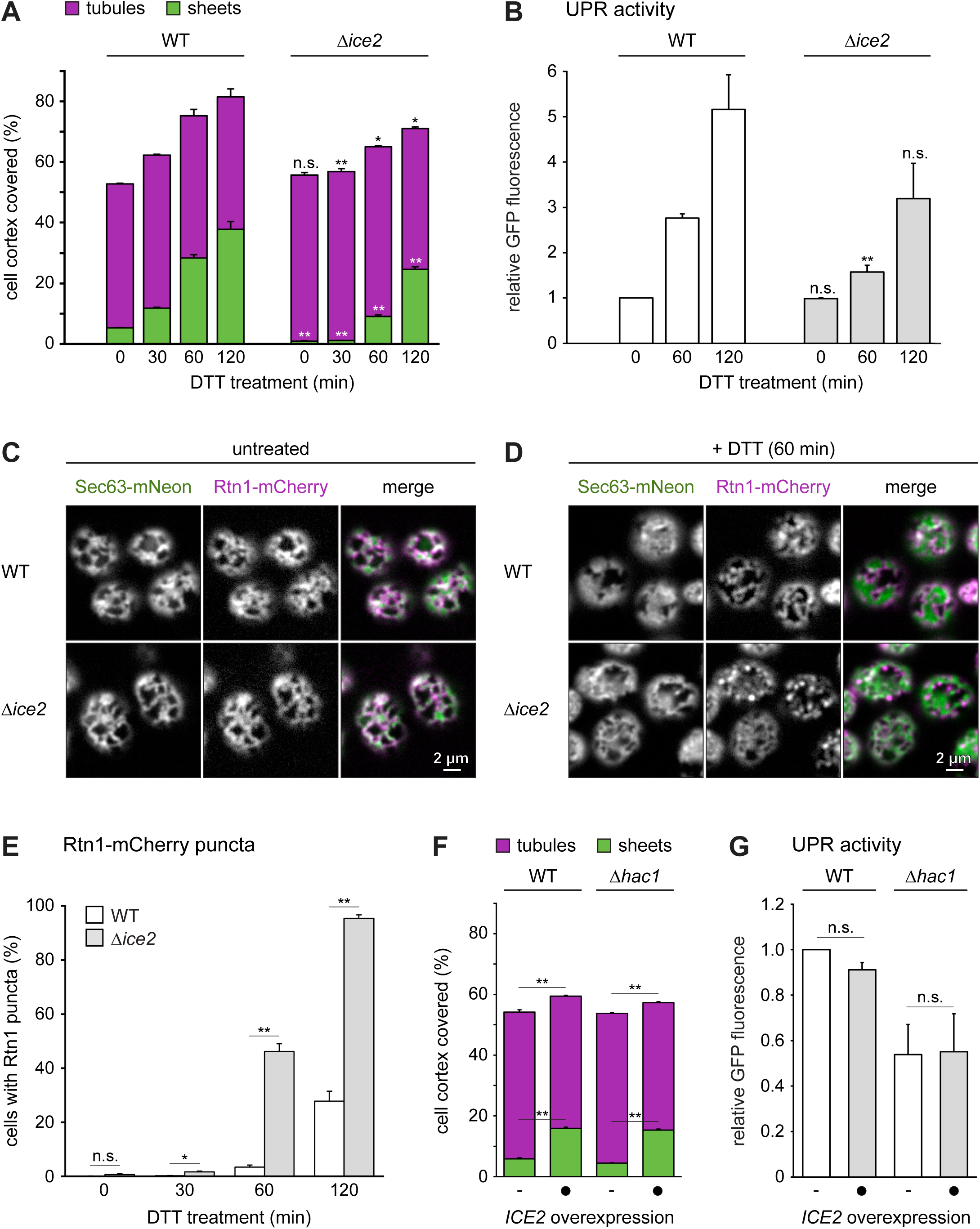
Ice2 is required for ER membrane biogenesis upon ER stress and *ICE2* overexpression is sufficient to induce ER expansion. **(A)** Quantification of peripheral ER structures in WT and *Δice2* cells (SSY1405, 1603) treated with 8 mM DTT for the times indicated. Plotted is the mean percentage of cell cortex covered by tubules (purple) or sheets (green), n = 3. Upper error bars are SEM for the sum of tubules and sheets, lower error bars are SEM for sheets. Asterisks indicate statistical significance compared with the corresponding value in wild-type cells. *, p < 0.05; **, p < 0.01; n.s., not significant. **(B)** Flow cytometric measurements of GFP levels of WT and *Δice2* cells containing the UPR reporter (SSY2306, 2312). Cells were treated with 8 mM DTT for the times indicated. Data were normalized to untreated WT cells. Mean ± SEM, n = 3. Asterisks indicate statistical significance compared with the corresponding value in WT cells. **(C)** Fluorescence images of cortical sections of untreated WT and *Δice2* cells expressing Sec63-mNeon and Rtn1-mCherry (SSY1405, 1603). **(D)** As in panel C, but after treatment with 8 mM DTT for 1 h. **(E)** Quantification of WT and *Δice2* cells with Rtn1-mCherry puncta after treatment with 8 mM DTT for the times indicated. Mean ± SEM. n = 3. **(F)** Quantification of peripheral ER structures in untreated WT and UPR-deficient *Δhac1* cells (SSY2228, 2331), overexpressing *ICE2* from plasmid pSS761 where indicated. Bars, error bars and asterisks are as in panel A. **(G)** Flow cytometric measurements of GFP levels of WT and *Δhac1* cells containing the UPR reporter (SSY2306, 2314) and overexpressing *ICE2* from plasmid pSS761 where indicated. Data were normalized to untreated WT cells. Mean ± SEM, n = 3.

Furthermore, induction of the UPR reporter by ER stress was reduced (Figure 4B). However, images of DTT-treated wild-type and *Δice2* cells revealed that ER expansion in *Δice2* mutants was not simply retarded but aberrant. Specifically, DTT treatment induced striking puncta positive for Rtn1-mCherry but not Sec63-mNeon, and these puncta were much more abundant in *Δice2* than in wild-type cells (Figure 4C-E). It remains to be determined whether these puncta are aberrant membrane structures or Rtn1-mCherry molecules not associated with the ER membrane. Either way, these data show that removal of Ice2 impairs ER expansion also during ER stress.

Finally, we asked whether raising Ice2 levels leads to ER expansion. Indeed, overexpression of *ICE2* caused ER expansion, and this still occurred in UPR-deficient *Δhac1* cells (Figure 4F; Emmerstorfer et al., 2015). In addition, *ICE2* overexpression did not activate the UPR (Figure 4G). Hence, Ice2 can drive ER membrane biogenesis independently of the UPR.

Collectively, these data show that Ice2 is required for and promotes ER membrane biogenesis. This impact of Ice2 does not result from disrupted Ino2/4 target gene induction in the absence of Ice2 or from UPR activation upon *ICE2* overexpression.

### Ice2 is functionally linked to Nem1, Spo7 and Pah1

Ice2 has been implicated in ER morphogenesis and lipid metabolism (Estrada de Martin et al., 2005; Tavassoli et al., 2013; Markgraf et al., 2014; Quon et al., 2018). In particular, Ice2 has been suggested to channel diacylglycerol (DAG) from lipid droplets (LDs) to the ER for phospholipid synthesis (Markgraf et al., 2014). We therefore asked whether defective ER membrane biogenesis in *Δice2* cells resulted from an insufficient supply of lipids from LDs. Deletion of *ICE2* impairs growth (Markgraf et al., 2014). Abolishing LD formation by combined deletion of *ARE1*, *ARE2*, *LRO1* and *DGA1* (Sandager et al., 2002) did not affect cell growth, and deletion of *ICE2* still impaired growth in the absence of LDs (Figure S1A). Therefore, Ice2 must have functions independent of LDs. Moreover, lack of LDs had no effect on ER expansion after ino2* expression or DTT treatment, and deletion of *ICE2* still impaired ER expansion in the absence of LDs (Figure S1B, C). Hence, the role of Ice2 in ER membrane biogenesis cannot be explained by LD-related functions. These results additionally show that ER expansion can occur without lipid mobilization from LDs.

Genome-scale studies have identified many genetic interactions of *ICE2* with lipid synthesis genes (Schuldiner et al., 2005; Costanzo et al., 2010; Surma et al., 2013). An interesting pattern emerged from mapping these data onto the biochemical pathways for membrane biogenesis and lipid storage (Figure 5A and Table S2). *ICE2* displays negative interactions with genes for membrane lipid synthesis but positive interactions with *NEM1*, *SPO7* and *PAH1*, which promote lipid storage. Positive genetic interactions, which cause the phenotype of a double mutant to be less severe than expected from the phenotypes of the respective single mutants, frequently reflect involvement of interacting genes in the same pathway (Costanzo et al., 2019). We therefore focused on the relationship of *ICE2* with *NEM1*, *SPO7* and *PAH1*.

**Figure 5.**
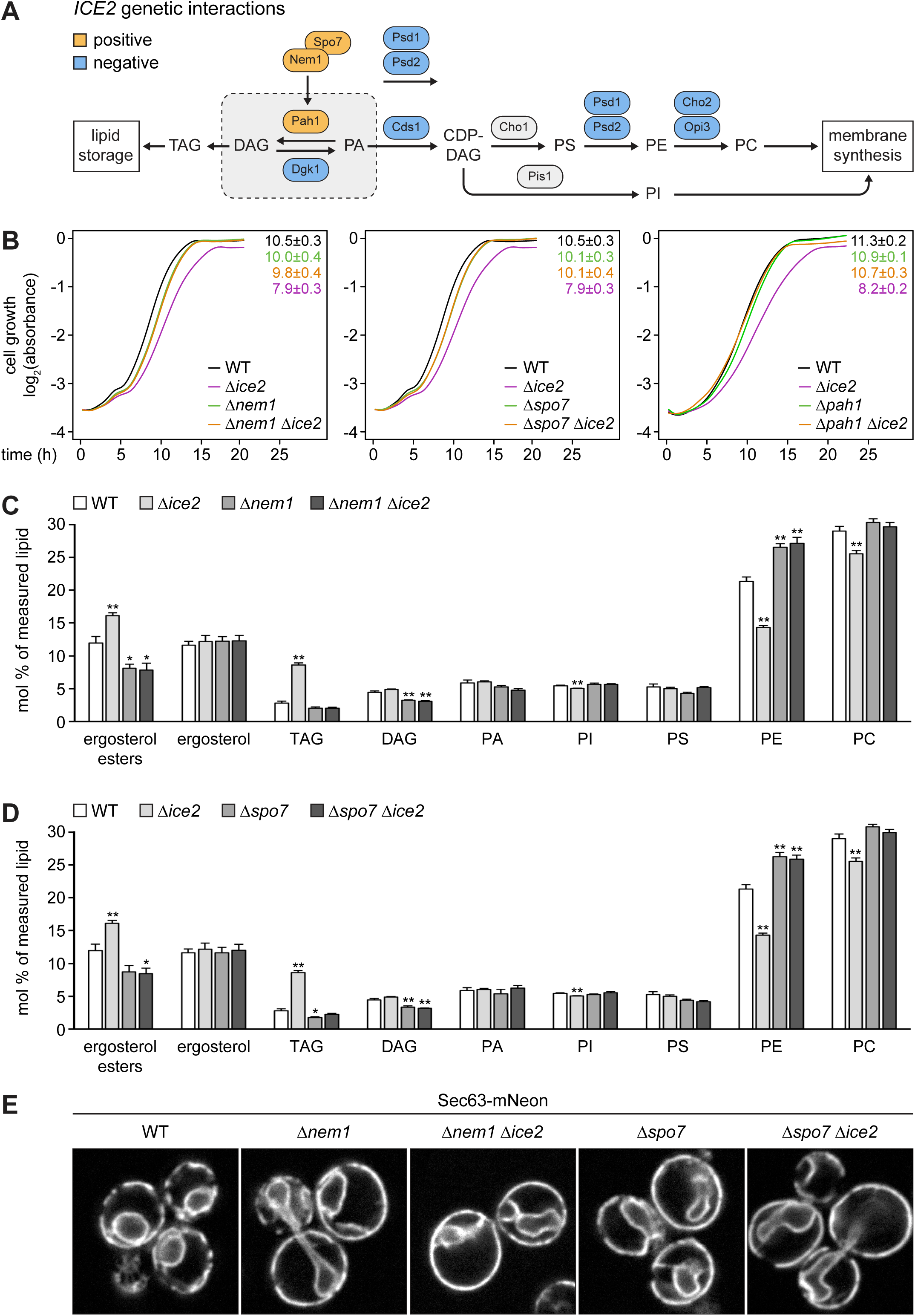
Ice2 is functionally linked to Nem1, Spo7 and Pah1. **(A)** Genetic interactions of *ICE2* with selected lipid synthesis genes. CDP, cytidine diphosphate; DAG, diacylglycerol; PA, phosphatidic acid; PI/PS/PE/PC, phosphatidyl- inositol/serine/ethanolamine/choline; TAG, triacylglycerol. **(B)** Growth assays of untreated WT, *Δice2*, *Δnem1*, *Δnem1 Δice2*, *Δspo7*, *Δspo7 Δice2*, *Δpah1* and *Δpah1 Δice2* cells (SSY1404, 2356, 2482, 2484, 2481, 2483, 2807, 2808). Numbers represent areas under the curves and serve as growth indices. Mean ± SEM, n = 3. Data for WT and *Δice2* cells are the same in the left and middle panels. **(C)** Lipidomic analysis of WT, *Δice2*, *Δnem1* and *Δice2 Δnem1* cells (SSY1404, 2356, 2482, 2484). Mean ± SEM, n = 4. Asterisks indicate statistical significance compared with the WT. *, p < 0.05; **, p < 0.01. **(D)** As in panel C, but of WT, *Δice2*, *Δspo7* and *Δice2 Δspo7* cells (SSY1404, 2356, 2481, 2483). Data for WT and *Δice2* cells are the same as in panel C. **(E)** Sec63-mNeon images of untreated WT, *Δnem1*, *Δnem1Δice2*, *Δspo7* and *Δspo7 Δice2* cells (SSY1404, 2482, 2484, 2481, 2483).

Pah1 is a phosphatidic acid phosphatase that converts PA into DAG (Figure 5A; Han et al. 2006). In yeast, PA is a precursor of all phospholipids, whereas DAG is the precursor of the storage lipid triacylglycerol (Henry et al., 2012). Pah1 thus promotes LD biogenesis (Adeyo et al., 2011). Pah1 is regulated by phosphorylation (Figure 6A). Phosphorylated Pah1 is cytosolic and inactive. Activation of Pah1 requires that it binds to and is dephosphorylated by the ER-localized Nem1-Spo7 complex, which consists of the phosphatase Nem1 and its indispensable binding partner Spo7 (Siniossoglou et al., 1998; Santos-Rosa et al., 2005; Karanasios et al., 2013). Dephosphorylated Pah1 associates with the ER and thus gains access to its substrate, PA (O’Hara et al., 2006; Karanasios et al., 2010). The Nem1-Spo7 complex therefore is an immediate upstream activator of Pah1.

**Figure 6.**
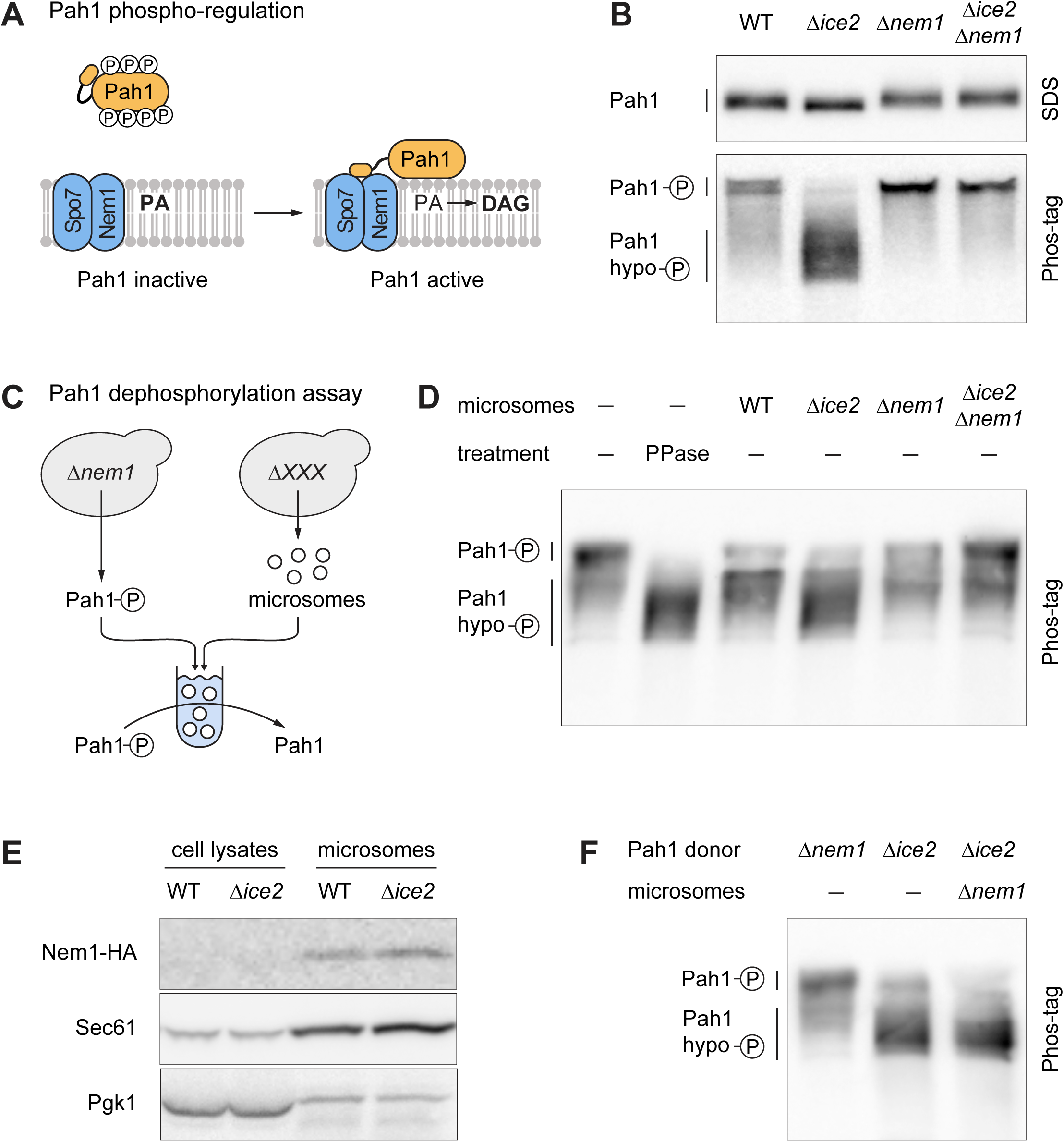
Ice2 opposes Pah1 by inhibiting the Nem1-Spo7 complex. **(A)** Schematic of Pah1 phospho-regulation. Phosphorylated Pah1 is cytosolic and inactive. Interaction of Pah1 and the ER-localized Nem1-Spo7 complex results in Pah1 dephosphorylation and activation, promoting conversion of PA into DAG. **(B)** Western blot of HA from WT, *Δice2*, *Δnem1* and *Δnem1 Δice2* cells expressing endogenously tagged Pah1- HA (SSY2592, 2593, 2594, 2718). SDS-PAGE and Phos-tag PAGE gels are shown. **(C)** Schematic of Pah1 dephosphorylation assay with phosphorylated Pah1 from *Δnem1* mutants and microsomes from different strains. **(D)** Western blot of HA from Pah1 dephosphorylation reactions that contained phosphorylated Pah1-HA from *Δnem1* mutants (SSY3065) and microsomes from cells of the indicated genotypes (SSY3053, 3074, 3075, 3095). Phosphorylated Pah1 and dephosphorylated Pah1 resulting from treatment with recombinant alkaline phosphatase (PPase) are shown for reference. **(E)** Western blot of HA, Sec61 and Pgk1 from cell lysates and microsomes prepared from WT and *Δice2* cells expressing Nem1-HA (SSY3140, 3141). Nem1 is undetectable in cell lysates due to its low abundance. **(F)** Western blot of HA from Pah1 phosphorylation reaction that contained hypophosphorylated Pah1- HA from *Δice2* mutants (SSY3096) and microsomes from *Δnem1* mutants (SSY3075). Phosphorylated Pah1 from *Δnem1* cells and dephosphorylated Pah1 from *Δice2* cells are shown for reference.

Growth assays confirmed the positive genetic interaction of *ICE2* with *NEM1*, *SPO7* and *PAH1*. Remarkably, the growth defect caused by *ICE2* deletion was completely prevented by deletion of any of the other three genes (Figure 5B). This type of strong positive genetic interaction is called suppression (Costanzo et al., 2019). One explanation for this relationship could be that Ice2 inhibits the Nem1-Spo7 complex and hence the activation of Pah1. In this scenario, removal of Ice2 would result in Pah1 overactivity, which is known to impair cell growth (Santos-Rosa et al., 2005), and additional removal of Nem1, Spo7 or Pah1 would suppress this phenotype. Removal of Ice2 would also result in an accumulation of triacylglycerol at the expense of phospholipids, and this effect would require the Nem1-Spo7 complex. We therefore analyzed the lipidomes of wild-type, *Δice2*, *Δnem1* and *Δspo7* cells. Compared with wild-type cells, *Δice2* mutants had increased levels of triacylglycerol and ergosterol esters, the two lipid classes that make up LDs, and decreased levels of phosphatidylethanolamine and phosphatidylcholine, the two major membrane phospholipids (Figure 5C, D). These changes agree with earlier data and confirm that deletion of *ICE2* enhances lipid storage (Markgraf et al., 2014). In contrast, the lipidomes of *Δnem1* and *Δspo7* cells showed changes in the opposite direction, confirming that deletion of *NEM1* or *SPO7* enhances membrane biogenesis (Siniossoglou et al., 1998). Furthermore, the lipidomes of *Δnem1 Δice2* and *Δspo7 Δice2* double mutants were indistinguishable from those of *Δnem1* and *Δspo7* single mutants, confirming that loss of *ICE2* is irrelevant in the absence of *NEM1* or *SPO7*. In agreement with the lipidomics data, deletion of *NEM1* or *SPO7* drastically expanded the ER and caused nuclear morphology defects, as reported (Siniossoglou et al., 1998; Campbell et al., 2006), and additional deletion of *ICE2* had no morphological effect (Figure 5E).

Overall, these results show that Ice2, Nem1, Spo7 and Pah1 are functionally linked. Furthermore, they support the idea that Ice2 acts as an inhibitor of Nem1, Spo7 and Pah1 in a pathway that controls lipid utilization and ER membrane biogenesis.

### Ice2 opposes Pah1 by inhibiting the Nem1-Spo7 complex

To test whether Ice2 is an inhibitor of the Nem1-Spo7-Pah1 pathway, we made use of the fact that the phosphorylation status of Pah1 is a well-established indicator of its activity (O’Hara et al., 2006). When we separated different phosphoforms of Pah1 on Phos-tag gels (Dubots et al., 2014), we found that removal of Ice2 caused dephosphorylation of Pah1, and this effect was dependent on Nem1 (Figure 6B).

These results show that Ice2 opposes Pah1 dephosphorylation, which it could achieve by inhibiting the Nem1-Spo7 complex. Alternatively, Ice2 could activate a Pah1 kinase. To distinguish between these possibilities, we reconstituted Pah1 dephosphorylation in vitro (Figure 6C). As substrate, we used hyperphosphorylated Pah1 immunoisolated from *Δnem1* cells. As source for the transmembrane proteins Nem1, Spo7 and Ice2, we used microsomes prepared from wild-type, *Δice2*, *Δnem1* or *Δice2 Δnem1* cells. Phosphorylated Pah1 ran as a low-mobility band on Phos-tag gels and shifted to higher mobility when dephosphorylated with alkaline phosphatase (Figure 6D). Incubation of hyperphosphorylated Pah1 with microsomes from wild-type cells caused partial Pah1 dephosphorylation. In comparison, microsomes from *Δice2* mutants yielded more extensive Pah1 dephosphorylation, whereas microsomes from *Δnem1* and *Δice2 Δnem1* mutants did not affect the phosphorylation status of Pah1. Hence, the phosphatase activity responsible for Pah1 dephosphorylation in this assay was provided by Nem1 and was stimulated by removal of Ice2. The levels of Nem1 in microsomes prepared from wild-type and *Δice2* cells were similar, ruling out that the high Nem1 activity in the absence of Ice2 resulted from increased Nem1 abundance (Figure 6E). Furthermore, we modified the in vitro assay to test whether the Pah1 phosphorylation status was affected by a kinase that could be activated by Ice2. Hypophosphorylated Pah1 immunoisolated from *Δice2* cells was incubated with microsomes from *Δnem1* cells so that any kinase activity targeting Pah1 could manifest itself without being masked by Nem1-mediated Pah1 dephosphorylation. No phosphorylation of Pah1 was apparent (Figure 6F), indicating that our assay exclusively reconstituted Pah1 dephosphorylation. Hence, Ice2 is an inhibitor of Nem1-mediated dephosphorylation of Pah1.

Ice2 could inhibit the Nem1-Spo7 complex by lowering its abundance. However, deletion of *ICE2* slightly destabilized Nem1 and Spo7, and *ICE2* overexpression had essentially no effect (Figure 7A, B). In contrast, deletion of *SPO7* strongly reduced the levels of Nem1, as reported (Mirheydari et al., 2020). Thus, it is likely that Ice2 negatively regulates the Nem1-Spo7 complex by inhibiting its phosphatase activity. To test whether Ice2 physically associates with the Nem1-Spo7 complex, we used co-immunoprecipitation. We chromosomally fused *NEM1* or *SPO7* with a FLAG tag and *ICE2* with an HA tag, solubilized the proteins with detergent and retrieved Nem1-FLAG or Spo7-FLAG with anti-FLAG antibodies. Ice2 co-precipitated with both Nem1 and Spo7 (Figure 7C, D). We were unable to test whether the association of Ice2 and Nem1 depends on Spo7 because Nem1 is unstable in the absence of Spo7 (Figure 7A; Mirheydari et al., 2020). However, Ice2 still co-precipitated with Spo7 in the absence of Nem1 (Figure 7E). Since Nem1 and Spo7 form a stable complex (Siniossoglou et al., 1998), these results suggest that Ice2, Nem1 and Spo7 can form a ternary complex. This notion, together with the fact that the Nem1-Spo7 complex physically interacts with Pah1 (Karanasios et al., 2013), implies that Ice2 is in the vicinity of a pool of Pah1. Indeed, fusion of Ice2 with the non-specific biotin ligase TurboID (Branon et al., 2018) resulted in biotinylation of Pah1. This proximity-dependent biotinylation was strongly reduced when ER recruitment of Pah1 was blocked by deletion of *NEM1* or *SPO7* (Figure 7F).

**Figure 7.**
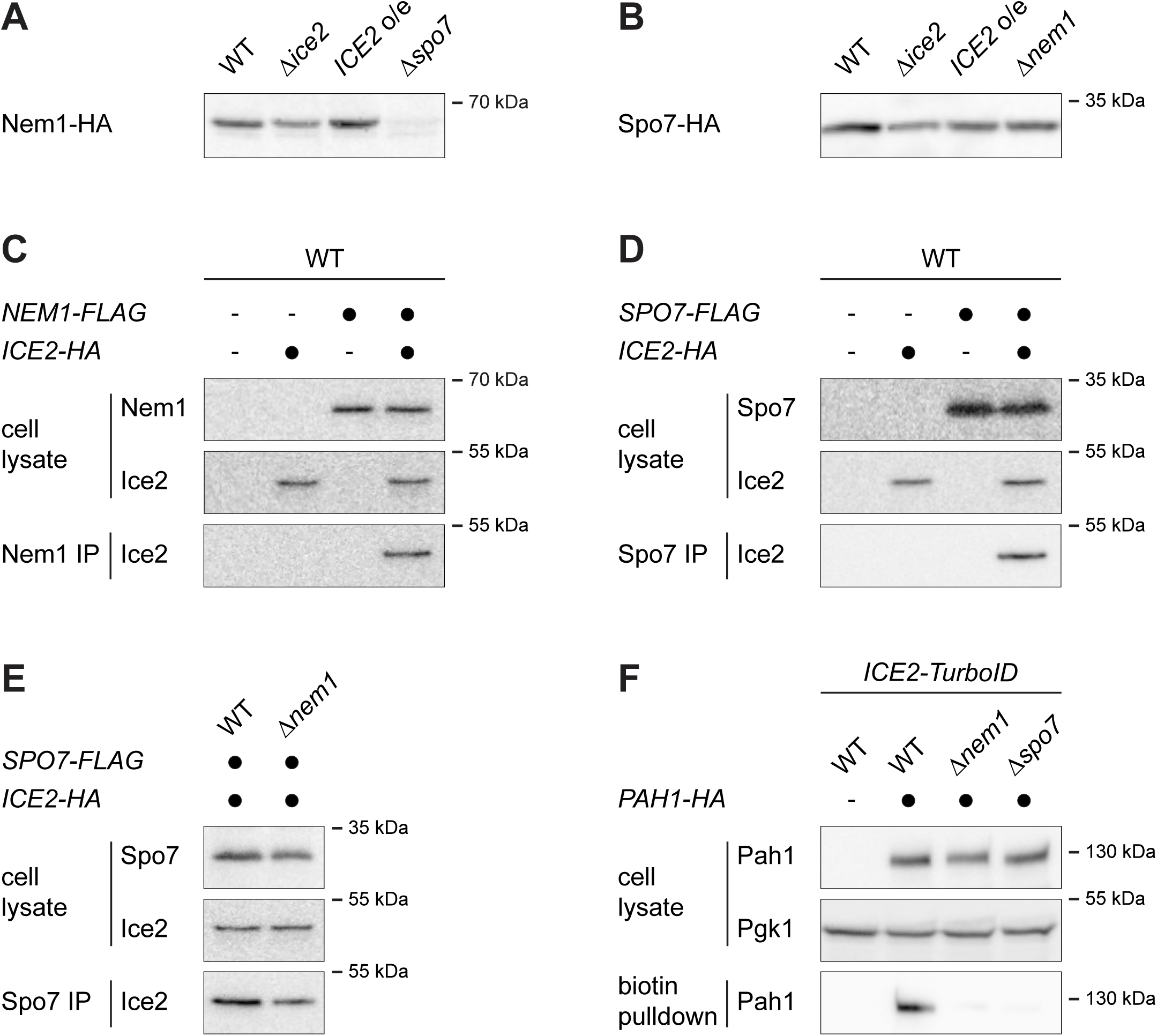
Ice2 forms a complex with Nem1 and Spo7. **(A)** Western blot of HA from total membranes prepared from WT, *Δice2*, *ICE2*-overexpressing and *Δspo7* cells containing Nem1-HA (SSY2913, 2914, 2915, 2945). Equal amounts of protein were loaded in each lane. **(B)** As in panel A but with WT, Δice2, *ICE2*-overexpressing and *Δnem1* cells containing Spo7-HA (SSY2910, 2911, 2912, 3182). **(C)** Western blots of HA and FLAG from cell lysates or anti-FLAG immunoprecipitates of cells containing Nem1-FLAG and Ice2-HA as indicated (SSY122, 2421, 3195, 3196). IP, immunoprecipitate. **(D)** As in panel C but with cells containing Spo7-FLAG and Ice2- HA as indicated (SSY122, 2421, 3183, 3184). **(E)** As in panel D but with WT and *Δnem1* cells containing Spo7-FLAG and Ice2-HA (SSY3184, 3197). **(F)** Western blot of HA and Pgk1 from cell lysates or biotin pulldowns of cells containing Ice2-TurboID, Pah1-HA and deletions of *NEM1* or *SPO7* as indicated (SSY2978, 2979, 3117, 3118).

Together, these findings show that Ice2 interacts with and inhibits the Nem1-Spo7 phosphatase complex, opposing dephosphorylation and thus activation of Pah1.

### Ice2 promotes ER membrane biogenesis through Pah1 phospho-regulation

We next tested whether Ice2 impacts ER biogenesis through inhibition of Pah1. We employed pah1(7A), which carries mutations in seven of the residues that are phosphorylated by various kinases and dephosphorylated by Nem1-Spo7 (O’Hara et al., 2006; Carman and Han, 2019). As a result, pah1(7A) is constitutively active, although some regulation by Nem1-Spo7 remains (Su et al., 2014). We quantified ER expansion upon *ICE2* overexpression in wild-type and pah1(7A)-expressing cells. Overexpression of *ICE2* in wild-type cells expanded the ER, as before (Figure 8A, also see Figure 4F). Replacement of Pah1 with pah1(7A) caused slight shrinkage of the ER at steady state, consistent with reduced membrane biogenesis. Moreover, pah1(7A) almost completely blocked ER expansion upon *ICE2* overexpression. Similarly, pah1(7A) impaired ER expansion upon DTT treatment, thus phenocopying the effects of *ICE2* deletion (Figure 8B, C, also see 4A, E). These data support the idea that Ice2 promotes ER membrane biogenesis by restricting Pah1 activity.

**Figure 8.**
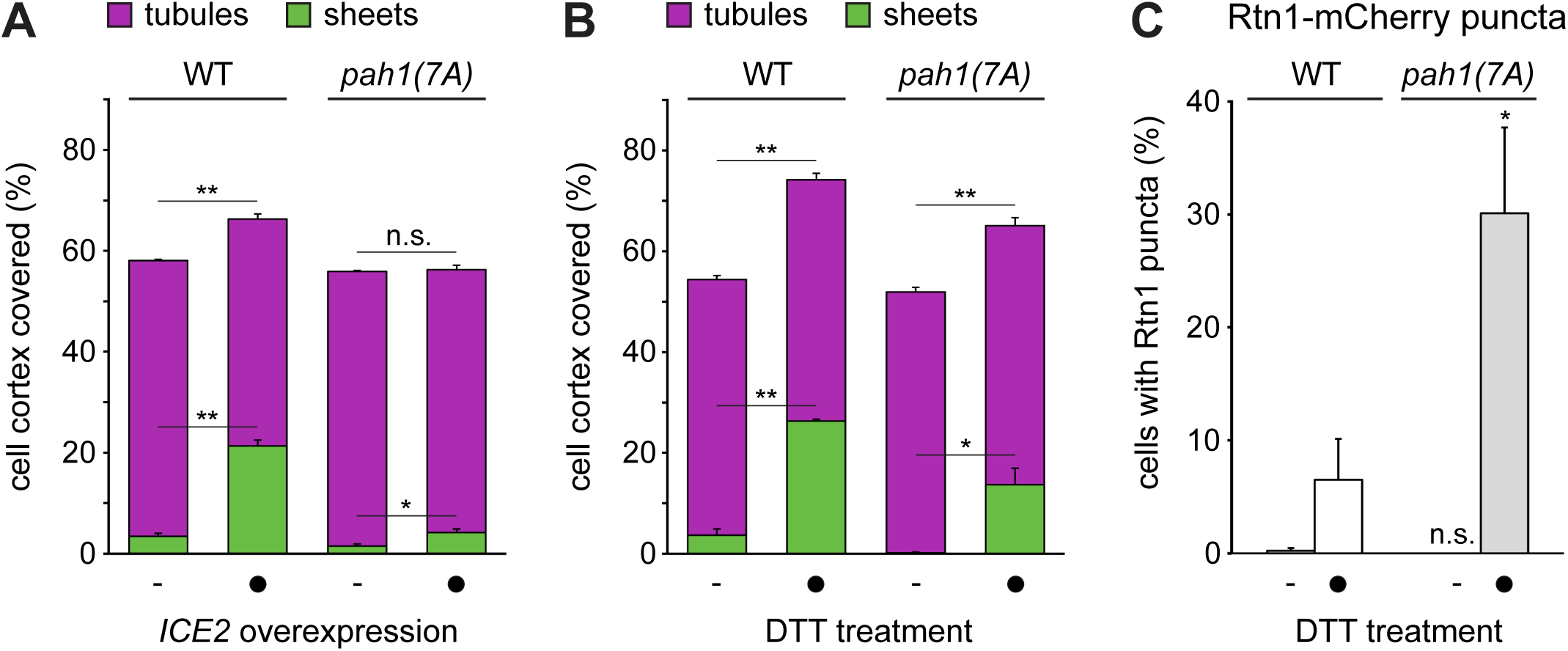
Ice2 promotes ER membrane biogenesis through Pah1 phospho-regulation. **A)** Quantification of peripheral ER structures in WT cells and cells in which *PAH1* was replaced by *pah1(7A)*, without and with overexpression of *ICE2* (SSY2841, 2842, 2843, 2844). Plotted is the mean percentage of cell cortex covered by tubules (purple) or sheets (green), n = 3. Upper error bars are SEM for the sum of tubules and sheets, lower error bars are SEM for sheets. Asterisks indicate statistical significance. *, p < 0.05; **, p < 0.01; n.s., not significant. **(B)** Quantification of peripheral ER structures in WT cells and cells in which *PAH1* was replaced by *pah1(7A)* (SSY2841, 2842), treated with 8 mM DTT for 1 h. Bars, error bars and asterisks as in panel A. **(C)** Quantification of WT and *Δice2* cells with Rtn1-mCherry puncta after treatment with 8 mM DTT for 1 h. Mean ± SEM, n = 3. Asterisks indicate statistical significance compared with the WT.

### Ice2 cooperates with the PA-Opi1-Ino2/4 system and promotes ER homeostasis

Given the important role of Opi1 in ER membrane biogenesis (Schuck et al., 2009), we asked how Ice2 is related to the PA-Opi1-Ino2/4 system. Deletion of *OPI1* and overexpression of *ICE2* both cause ER expansion. These effects could be independent of each other or they could be linked. Combined deletion of *OPI1* and overexpression of *ICE2* produced an extreme ER expansion, which exceeded that in *Δopi1* mutants or *ICE2*-overexpressing cells (Figure 9A, B). This hyperexpanded ER covered most of the cell cortex and contained an even greater proportion of sheets than the ER in DTT-treated wild-type cells (Figure 9B, also see Figure 4A). Therefore, Ice2 and the PA-Opi1-Ino2/4 system make independent contributions to ER membrane biogenesis.

**Figure 9.**
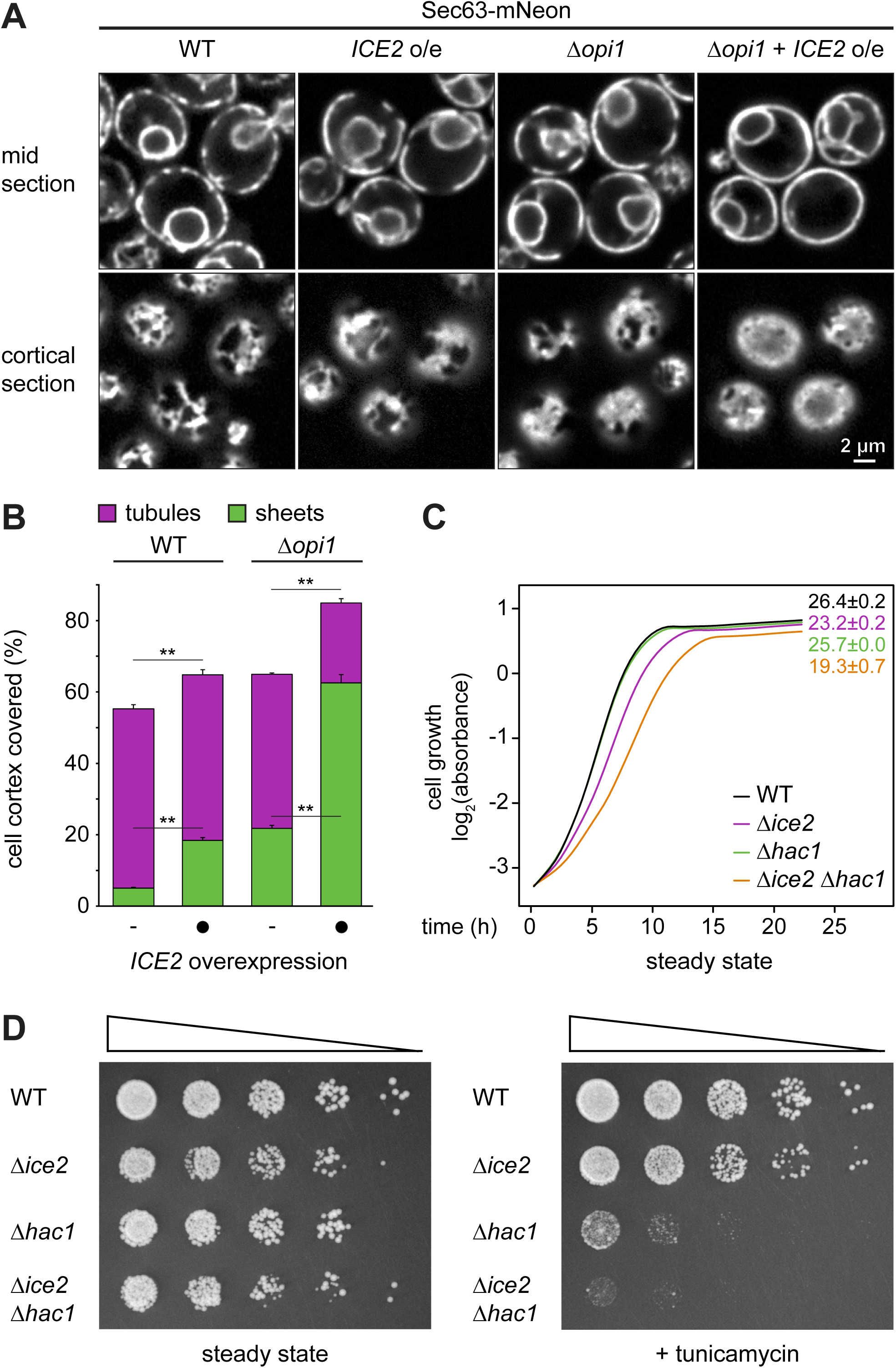
Ice2 cooperates with the Opi1-Ino2/4 system and promotes ER homeostasis. **(A)** Sec63-mNeon images of mid and cortical sections of untreated WT and *Δopi1* cells, overexpressing *ICE2* where indicated (SSY1404, 2588, 2595, 2596). **B)** Quantification of peripheral ER structures in the strains shown in panel A. Plotted is the mean percentage of cell cortex covered by tubules (purple) or sheets (green), n = 3. Upper error bars are SEM for the sum of tubules and sheets, lower error bars are SEM for sheets. Asterisks indicate statistical significance. **, p < 0.01. **(C)** Growth assays of untreated WT, *Δhac1*, *Δice2*, and *Δhac1 Δice2* cells (SSY1404, 2356, 2805, 2806). Numbers represent areas under the curves and serve as growth indices. Mean ± SEM, n = 3. **(D)** Growth assays on solid media of WT, *Δhac1*, *Δice2*, and *Δhac1 Δice2* cells (SSY1404, 2356, 2805, 2806) in the absence or presence of 0.2 µg/ml tunicamycin. For each series, cells were diluted fivefold from one step to the next.

Last, to gain insight into the physiological significance of Ice2, we analyzed the interplay of Ice2 and the UPR. Under standard culture conditions, *Δice2* mutants show a modest growth defect (Markgraf et al., 2014), and UPR-deficient *Δhac1* mutants grow like wild-type cells (Sidrauski et al., 1996). Nevertheless, *Δice2 Δhac1* double mutants grew slower than *Δice2* mutants (Figure 9C). This synthetic phenotype was even more pronounced under ER stress. In the presence of the ER stressor tunicamycin, which we prefer over DTT for growth assays because of its stability, *Δice2* mutants showed a slight growth defect, *Δhac1* mutants showed a strong growth defect, and *Δice2 Δhac1* double mutants showed almost no growth at all (Figure 9D). Hence, Ice2 is particularly important for cell growth when ER stress is not buffered by the UPR. These results emphasize that Ice2 promotes ER homeostasis.

## DISCUSSION

In this study, we have identified factors involved in ER membrane expansion upon enforced lipid synthesis in yeast. We show that Ice2 is critical for proper ER expansion, both upon enforced lipid synthesis and during ER stress. We find that Ice2 inhibits the Nem1-Spo7 complex, thereby opposing the activation of the phosphatidic acid phosphatase Pah1 and promoting membrane biogenesis. These observations uncover an additional layer of regulation of the Nem1-Spo7-Pah1 phosphatase cascade. Finally, we provide evidence that Ice2 cooperates with the PA-Opi1-Ino2/4 system to regulate ER membrane biogenesis and helps to maintain ER homeostasis.

Our findings can be integrated into a model of the regulatory network that controls ER membrane biogenesis (Figure 10). At the core of this network is the interconversion of DAG and PA by Dgk1 and Pah1. Ice2 inhibits Pah1 dephosphorylation by the Nem1- Spo7 complex and thus suppresses conversion of PA into DAG. The resulting increased availability of PA is coordinated with the production of lipid synthesis enzymes that turn PA into other phospholipids. Specifically, inhibiting Pah1 prevents it from repressing Ino2/4-controlled lipid synthesis genes (Santos-Rosa et al., 2005). In addition, PA sequesters Opi1 and thereby derepresses Ino2/4 target genes (Loewen et al., 2004). Hence, inhibition of Pah1 by Ice2 increases the availability of PA and, concomitantly, induces phospholipid synthesis genes.

**Figure 10.**
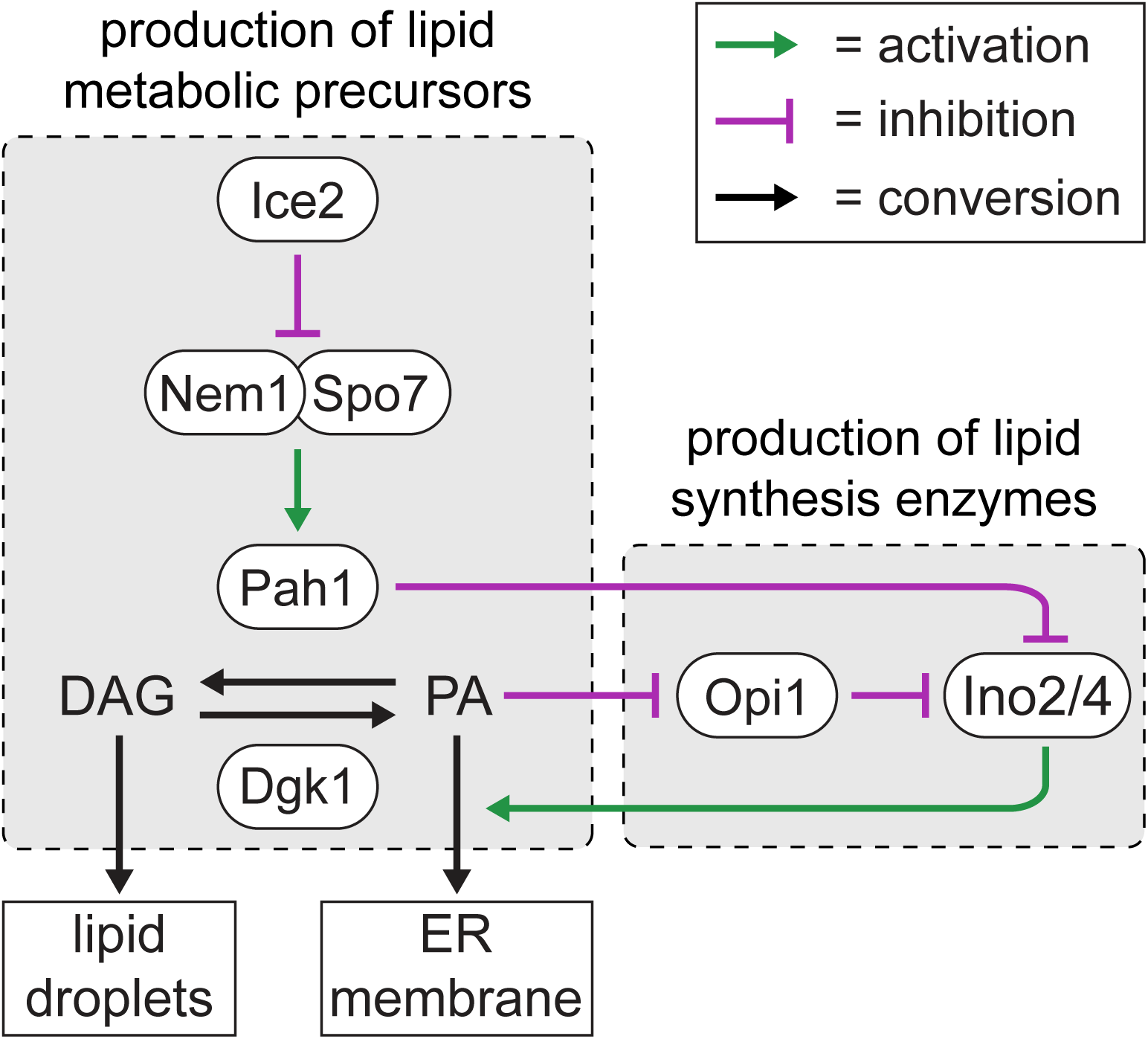
Model for the regulation of ER membrane biogenesis. Pah1 converts phosphatidic acid (PA) into diacylglycerol (DAG) for lipid droplet biogenesis. Ice2 inhibits the Nem1-Spo7 complex and hence Pah1. Ice2 thereby increases PA availability and relieves repression of Ino2/4-driven lipid synthesis genes, thus promoting ER membrane biogenesis. These mechanisms coordinate the production of lipid metabolic precursors and lipid synthesis enzymes.

This model readily explains the effects of *ICE2* deletion and overexpression. The disruption of ino2*-driven ER expansion by *ICE2* deletion may be explained by the need to coordinate the production of lipid metabolic precursors with the expression of lipid synthesis genes. As we show, ino2* still induces genes encoding lipid synthesis enzymes in *Δice2* mutants. Nevertheless, ER expansion fails, likely because the supply of substrates for these enzymes is insufficient. The same reasoning can explain the additive effects of *OPI1* deletion and *ICE2* overexpression on ER membrane biogenesis. *ICE2* overexpression increases PA availability and presumably releases some repression of Ino2/4 target genes. However, *OPI1* deletion likely further boosts the levels of lipid synthesis enzymes. Conversely, constitutive Pah1 activation upon *ICE2* deletion can explain the increase in LD abundance in *Δice2* mutants (Markgraf et al., 2014), although our data do not exclude direct roles of Ice2 in lipid mobilization from LDs. An important topic for future investigation is the regulation of Ice2. The *ICE2* gene is not induced by ER stress (Pincus et al., 2014). A possibility to be explored is that Ice2 activity is controlled by phosphorylation, as is the case not only for Pah1 but also for Nem1 and Dgk1 (Kwiatek et al., 2020).

How could our findings from yeast apply to higher eukaryotes? Bioinformatic analysis suggests mammalian SERINC proteins as distant Ice2 orthologs (Alli-Balogun and Levine, 2021). SERINC1 was initially proposed to be at the ER, but later work detected SERINC proteins only at endosomes and the plasma membrane (Inuzuka et al., 2005; Rosa et al., 2015; Davies et al., 2018). Whether SERINC proteins have similar roles as Ice2 remains to be tested. In contrast, Nem1, Spo7 and Pah1 are evolutionarily conserved (Han et al., 2012). The mammalian Pah1 orthologs lipin-1/2/3 are phospho- regulated in a similar manner as Pah1 and function in lipid storage in mice and humans (Harris and Fink, 2011; Zhang and Reue, 2017). Like in yeast, removal of lipins in protozoa, plants, worms and flies causes ER expansion (Golden et al., 2009; Eastmond et al., 2010; Bahmanyar et al., 2014; Grillet et al., 2016; Pillai et al., 2017). However, while both yeast and metazoa use the CDP-DAG pathway to synthesize phosphatidylinositol, metazoa generate the major phospholipids phosphatidylcholine and phosphatidylethanolamine mainly through the Kennedy pathway (Vance, 2015). The Kennedy pathway uses DAG as a precursor for phospholipids and not PA. Therefore, DAG is a precursor for both LD and membrane biogenesis, seemingly excluding the possibility that the balance between DAG and PA could determine whether LD or ER biogenesis is favored. This incongruence may be resolved by the finding in *A. thaliana* that CCT, the rate-limiting enzyme for phosphatidylcholine synthesis by the Kennedy pathway, is allosterically activated by PA (Craddock et al., 2015). Activation of CCT by PA may also exist in mice (Zhang et al., 2019). Thus, accumulation of PA may increase flux through the Kennedy pathway and channel DAG towards phospholipid synthesis (Jacquemyn et al., 2017). This model, while speculative, raises the unifying possibility that lipins inversely govern LD and ER biogenesis in all eukaryotes.

Lipins act at the ER but are also found at lipid droplets, mitochondria, endosomes and inside the nucleus (Zhang and Reue, 2017). It appears plausible that different organelles use distinct mechanisms to recruit lipins and possess different regulators of lipin activity. Interestingly, we find that *ICE2* overexpression expands the peripheral ER but does not obviously alter the morphology of the nucleus (Figure 9A). This is in contrast to deletion of *PAH1*, which leads to both peripheral ER expansion and nuclear membrane proliferation (Santos-Rosa et al., 2005). Given that there are distinct intra- and extranuclear pools of Pah1 (Romanauska and Köhler, 2018), it appears possible that the intranuclear pool of Pah1 is responsible for maintaining proper nuclear morphology and is controlled in a way that does not involve Ice2. Furthermore, recent work in flies showed that the AAA-type ATPase Torsin specifically inhibits nuclear lipin by removing the Nem1 ortholog CTDNEP1 from the nuclear envelope (Jacquemyn et al., 2020). Thus, organelle-specific regulators of lipins may be important determinants of local lipin activity.

The regulation of lipid metabolism determines whether lipids are consumed, stored or used to build membranes. Elucidating how this decision is made will yield a deeper understanding of differentiation processes, for example the massive ER expansion during plasma cell development or the huge increase in lipid droplet abundance during adipogenesis. Moreover, it may enable therapeutic intervention in diseases associated with ER overload and aberrant lipid metabolism, such as diabetes and obesity. Besides organelle biogenesis, lipin activity impacts autophagy, axon regeneration, myopathy, dystonia and neurodegeneration (Zhang et al., 2014; Grillet et al., 2016; Zhang and Reue, 2017; Fanning et al., 2019; Yang et al., 2019; Schäfer et al., 2020). Unraveling the regulation of lipin will therefore have implications for a large variety of cellular processes and associated diseases.

## MATERIALS AND METHODS

### Plasmids

Plasmids used in this study are listed in Table S3. To generate pNH605-P_ADH1_-GEM- P_GAL1_-ino2(L119A), the *ino2(L119A)* sequence was subcloned from pRS415_MET25_- ino2(L119A) into pRS416-P_GAL1_. The P_GAL1_-ino2(L119A)-T_CYC1_ cassette was then transferred into pNH605-P_ADH1_-GEM. Similarly, pRS306-P_ADH1_-GEM-P_­_- ino2(L119A) was generated by transferring the P_GAL1_-ino2(L119A) cassette from pRS416-P_GAL1_-ino2(L119A) into pRS306-P_ADH1_-GEM. To generate pFA6a-mNeon- kanMX6 and pFA6a-mNeon-HIS3MX6, mNeonGreen (Shaner et al., 2013) was amplified from pFA6a-mNeon-kanMX4 and inserted into pFA6a-GFP(S65T)-kanMX6 and pFA6a-GFP(S65T)-HISMX6, replacing GFP(S65T). To generate pRS415-P_ADH1_- Ice2, the *ICE2* coding sequence was amplified from yeast W303 genomic DNA and inserted into pRS415-P_ADH1_. To generate plasmids encoding HA-tagged Pah1, YCplac111-Pah1-PrtA was linearized by inverse PCR and re-ligated using the NEBuilder HiFi DNA assembly mix (New England Biolabs, Ipswitch, Massachusetts) so that the Protein A sequence was replaced by a triple HA sequence. The segment of the *pah1(7A)* sequence containing the seven alanine substitutions was amplified from YCPlac111-pah1(7A)-PrtA and inserted into YCPlac111-Pah1-3HA to yield YCplac111-pah1(7A)-3HA.

### Yeast strain generation and growth

Strains used in this study are listed in Table S4. Strains generated for screening procedures were in the S288C background. All other strains were in the W303 background. Chromosomal modifications were introduced using PCR products or linearized plasmids (Longtine et al., 1998; Janke et al., 2004). The marker-free strains SSY2836 and SSY2837 were derived from SSY2809 by transformation with the Pah1- 3HA or the pah1(7A)-3HA sequence amplified from plasmids pSS1045 or pSS1047 followed by counterselection on 5-fluoroorotic acid.

Strains were grown at 30°C on YPD, SCD-MSG or SCD medium as indicated. YPD medium consisted of 1% yeast extract (Becton Dickinson, Heidelberg, Germany), 2% peptone (Becton Dickinson) and 2% glucose (Merck, Darmstadt, Germany). SCD-MSG medium consisted of 0.17% yeast nitrogen base without amino acids and ammonium sulfate (Formedium, Norfolk, UK), 0.1% monosodium glutamate, amino acids and 2% glucose. SCD medium consisted of 0.7% yeast nitrogen base without amino acids (Sigma, Taufkirchen, Germany), amino acids and 2% glucose. Media contained antibiotics or lacked certain amino acids as appropriate for selection.

### Light microscopy

Precultures were grown in liquid SCD medium during the day, diluted into fresh medium and grown overnight for 16 h so that they reached mid log phase (OD_600_ = 0.5 - 1). For induction of the ER biogenesis system, overnight cultures were diluted to OD_600_ = 0.05 in fresh medium and treated with the indicated concentrations of ß- estradiol (Sigma) for up to 6 h. For DTT treatment, overnight cultures were diluted to OD_600_ = 0.1 and treated with 8 mM DTT (Roche, Mannheim, Germany) for up to 2 h. Immediately before imaging, cells were harvested by centrifugation, mounted on coverslips and covered with a 1% (w/v) agarose pad. Images were acquired with a DMi8 inverted microscope (Leica, Wetzlar, Germany) equipped with a CSU-X1 spinning-disk confocal scanning unit (Yokogawa, Musashino, Japan) and an ORCA-Flash 4.0 LT camera (Hamamatsu, Hamamatsu, Japan). A HC PL APO 63x/1.40-0.60 or a HC PL APO 100x/1.4 CS2 oil objective lens (Leica) was used.

### UPR assays

UPR activity was measured by flow cytometry as described (Schmidt et al., 2019). Cells expressing cytosolic BFP and the 4xUPRE-GFP transcriptional UPR reporter (Jonikas et al., 2009) were grown to mid log phase and treated with estradiol or DTT as above. Fluorescence was measured with a FACS Canto flow cytometer (BD Biosciences, Franklin Lakes, New Jersey) equipped with a high-throughput sampler. Background autofluorescence was determined with identically grown isogenic control strains not harboring the UPR reporter. Background-subtracted GFP fluorescence was divided by BFP fluorescence to account for differences in protein translation capacity. GFP/BFP ratios were normalized to untreated wild-type cells.

### Cell lysis and western blotting

For standard western blotting, cells were harvested by centrifugation, washed once with water, resuspended in 50 mM HEPES pH 7.5 containing 0.5 mM EDTA, 1 mM PMSF and protease inhibitors (cOmplete, Roche), and disrupted by bead beating with a FastPrep 24 (MP Biomedicals). SDS was added to 1.5% (w/v) and proteins were solubilized by incubation at 65°C for 5 min. Equal protein amounts, as determined with the BCA assay kit (Thermo Fisher Scientific, Waltham, Massachusetts), were resolved by SDS-PAGE and transferred onto nitrocellulose membranes. Membranes were probed with primary and HRP-coupled secondary antibodies and developed with homemade ECL or SuperSignal West Femto maximum sensitivity substrate (Thermo Fisher). Chemiluminescence was detected with an ImageQuant LAS 4000 imaging system (GE Healthcare, Chalfont St Giles, UK). Primary antibodies were rabbit anti- Sec63 (Feldheim et al., 1992), mouse anti-mCherry 1C51 (Abcam, Cambridge, UK), rabbit anti-Sec61 (Schuck et al., 2009), mouse anti-Pgk1 22C5 (Abcam), rat anti-HA 3F10 (Roche) and mouse anti-FLAG M2 (Sigma).

For Phos-tag PAGE of total cell lysates, cells were lysed by alkaline extraction and incubation in a modified SDS sample buffer at 65°C for 3 min (Kushnirov, 2000). Proteins were resolved on zinc-containing Phos-tag gels according to Nagy et al., 2018, with minor modifications. Gels contained 8% acrylamide/bisacrylamide (29:1), 25 µM Phos-tag acrylamide (Fujifilm Wako Chemicals, Neuss, Germany) and 50 µM ZnCl_2_ in 350 mM Bis-Tris-HCl pH 6.8 and were run at 200 V at room temperature. An alternative, but equivalent, protocol was used for samples from in vitro assays. Samples were combined with 4x regular SDS sample buffer to adjust sample buffer concentration to 1x, incubated at 65°C for 5 min and resolved on 8% gels containing 50 µM Phos-tag acrylamide and 100 µM MnCl_2_ in 400 mM Tris-HCl pH 8.8. These gels were run at 80 V at 4°C for 20 min, followed by running at 15 mA/gel. To remove metal ions, gels were washed 3 x 10 min with transfer buffer (25 mM Tris-HCl, 192 mM glycine, 20% ethanol) containing 1 mM EDTA and 2 x 20 min with transfer buffer. Blotting was done at 100 V at 4°C for 3 h and blots were developed as above.

### Construction of the diploid ER marker knockout library

Using strains SSY2589 and SSY2590, the ER marker proteins Sec63-mNeon and Rtn1-mCherry and the GEM-P_GAL1_-ino2* cassette were integrated into a yeast knockout collection through modified SGA methodology (Giaever et al., 2002; Tong and Boone 2007). SSY2589 and SSY2590 were independently mated to the knockout collection on YPD medium using a Singer RoToR robot. For each library, diploids were selected on SCD-MSG lacking uracil and containing G418. Cells were pinned onto enriched sporulation medium (1% potassium acetate, 0.1% yeast extract, 0.05% glucose, 0.01% amino acid supplement consisting of only histidine, leucine, lysine and uracil) and kept at 23°C for 5 days. Haploids were selected by two rounds of pinning onto SCD-MSG lacking histidine/arginine/lysine with canavanine and thialysine or SCD-MSG lacking leucine/arginine/lysine with canavanine and thialysine to select for MATa (Sec63-mNeon library) and MATα (Rtn1-mCherry library) cells, respectively. Haploid cells harboring the required markers were selected by sequential pinning onto appropriate media. The two libraries were then mated together and diploids were selected on YPD containing nourseothricin and hygromycin to generate the final library with the genotype: *xxxΔ::kan/xxxΔ::kan SEC63-mNeon::HIS3/SEC63 RTN1- mCherry::nat/RTN1 can1Δ::GEM-P_GAL1_-ino2*-URA3/can1Δ::P_STE2_-HIS3 lyp1Δ::GEM- P_GAL1_-ino2*-URA3/lyp1Δ::P_STE3_-LEU2 his3Δ::P_GPD_-TagBFP-hph/his3Δ0.* This diploid library afforded two benefits compared with a haploid library. The larger size of diploid cells facilitated acquisition of informative images and the fact that cells were heterozygous for the fluorescently tagged ER marker proteins reduced the risk that specious phenotypes arose from impaired Sec63 or Rtn1 function.

### Automated microscopy

Cells were grown to saturation overnight in 100 *µ*l SCD medium in regular 96 well microtiter plates. Prior to imaging, 7 *µ*l of culture was transferred into 1 ml fresh SCD medium containing 800 nM estradiol and grown for 5 h in 96 deep-well microtiter plates to reach logarithmic growth phase. One-hundred microliters of each sample was transferred into 96 well glass bottomed microtiter plates (Brooks Life Sciences, Chelmsford, Massachusetts) coated with concanavalin A and allowed to attach. Medium was refreshed after 1 h to remove non-attached cells. Samples were imaged with a Nikon Ti-E wide-field microscope equipped with a motorized stage, a Nikon perfect focus system, a Flash4 Hamamatsu sCMOS camera and a 60x/1.49 oil immersion lens. For each sample, two fields of view were acquired consisting of five optical slices spaced 1 µm apart. Untreated wild-type control strains were included in duplicate on each plate as a reference for unexpanded ER.

### Automated cell segmentation and ER size measurement

Image analysis was done in MATLAB using custom scripts. Initial cell objects were identified based on the cytoplasmic BFP. The best overall mid-section image was selected by assessing the standard deviation in the BFP image, which is highest when the image is in focus. Next, fast Fourier transformation and bandpass filter were used to enhance contrast at the cell border. A Frangi filter (based on the implementation by D.J. Kroon, “Hessian based Frangi Vesselness filter”, MATLAB Central File Exchange. Retrieved May 2017) followed by Otsu thresholding was then used to generate a mask of apparent cell borders. Morphological opening followed by a minimum size filter was used to remove false labeling created by yeast vacuoles. The resulting image highlighted the cell borders. Because the borders of many cells touched each other, the internal space was used to identify and separate the individual cell objects. The intensity of these initial cell objects was measured and only objects brighter than two median absolute deviations below the median were kept. Finally, any remaining touching cells, including connected mother cells and buds, were separated by watershedding. The segmentation of individual cell objects thus obtained was then optimized to generate more accurate cell boundaries and peripheral ER segmentation. For this, cells were cropped and the best mid-section was reassessed on a per cell basis using the standard deviation of the Rtn1-mCherry image. The BFP images were re-segmented using the above procedure based on the new mid-section. To accurately define the cell periphery for image quantification, object borders were expanded but contained within watershed boundaries. The ER was segmented in both the Sec63-mNeon and Rtn1-mCherry images using a Frangi tubeness filter. A more accurate cell border was defined by fitting a minimum volume ellipse (based on the implementation by N. Moshtagh, “Minimum Volume Enclosing Ellipsoid”, MATLAB Central File Exchange. Retrieved July 2017) to the combined masks of the segmented ER. Based on this segmentation, cell area, mean Sec63-mNeon and Rtn1-mCherry fluorescence, and cell roundness were calculated. An area five pixels from the border was used to define the cell periphery area. Segmented ER falling within this area was used to define the peripheral ER area. From this, peripheral ER size (peripheral ER area divided by cell periphery area), ER profile size (mean area of ER profiles divided by cell periphery area) and number of ER gaps (number of gaps in the peripheral ER mask per micrometer cell periphery length) were calculated. Finally, to remove false cell objects, poorly segmented cells and dead cells, all of these measurements were used to limit the cell population to values within 2.5 standard deviations of the population mean. On average, 248 cells were analyzed per mutant, with the minimum being 25.

### Visual ER morphology analysis

Images were assessed visually using a custom image viewer application made in MATLAB. Segmented cells were arrayed in montages displaying 7 x 15 cells at a time. ER morphologies were independently annotated by two individuals with one or more of the following features: underexpanded, overexpanded, extended sheets, disorganized and clustered. All strains with abnormal ER morphology were re-imaged to ensure that the phenotype was robust.

### Computational ER expansion analysis

Since most gene deletions did not affect ER expansion, mutants from the same imaging plate served as a plate-specific background population for comparison to individual deletion strains. Sec63-mNeon intensity was used to define this background population and exclude the influence of extreme outliers. First, to remove plate effects, mNeon intensities were normalized by subtracting the plate means. Next, values were corrected for cell size (bigger cells being brighter) and cell count (densely crowded areas having an overall higher fluorescence) by local regression. Finally, the background population (BP) was defined for each plate as mutants that were within 1.5 standard deviations of the mean. To normalize the ER expansion measurements, a Z score was calculated as (sample - BP mean)/BP standard deviation, thereby removing plate effects. The time spent imaging each plate (approximately 50 minutes) was accounted for by correcting for well order by local regression. Similarly, cell density effects were corrected for by local regression against cell count. Scores were calculated separately for each field of view and the maximum value was taken for each sample. False positives were removed by visual inspection, which were often caused by an out of focus field of view. Strains passing arbitrary thresholds of significance (Z score less than -2 for total peripheral ER size and ER profile size, and greater than 2 for ER gaps) in at least two of the measurements and no overall morphology defects as defined above were re-imaged in triplicate along with wild-type control strains under both untreated and estradiol-treated conditions. Images were inspected visually as a last filter to define the final list of strains with ER expansion defects.

### Semi-automated cortical ER morphology quantification

For cell segmentation, bright field images were processed in Fiji to enhance the contrast of the cell periphery. For this, a Gaussian blur (sigma = 2) was applied to reduce image noise, followed by a scaling down of the image (x = y = 0.5) to reduce the effect of small details on cell segmentation. A tubeness filter (sigma = 1) was used to highlight cell borders and the images were scaled back up to the original resolution. Cells were segmented using CellX (Dimopoulos et al., 2014) and out of focus cells were removed manually. A user interface in MATLAB was then used to assist ER segmentation (available at https://github.com/SchuckLab/ClassifiER). The user inputs images of Sec63-mNeon and Rtn1-mCherry from cortical sections (background subtracted in Fiji using the rolling ball method with a radius of 50 pixels) and the cell segmentation file generated in CellX. Adjustable parameters controlled the segmentation of ER tubules and sheets for each image. These parameters were tubule/sheet radius, strength and background. Manual fine-tuning of these parameters was important to ensure consistent ER segmentation across images with different signal intensities. These parameters were set independently for Sec63-mNeon and Rtn1-mCherry images together with one additional parameter called ‘trimming factor’, which controls the detection of ER sheets. ER masks were calculated across entire images and were assigned to individual cells based on the CellX segmentation. For each channel, the background (BG) levels were automatically calculated using Otsu thresholding and fine-tuned by multiplying the threshold value by the ‘tubule BG’ (Rtn1 channel) or ‘total ER BG’ (Sec63 channel) adjustment parameters. A 3x3 median filter was applied to smoothen the images and reduce noise that is problematic for segmentation. Two rounds of segmentation were passed for each image channel (Sec63 or Rtn1) with one optimized for finding smaller features (tubules) and the other for larger features (sheets). Firstly, convolution kernels were calculated for small and large features, respectively, defined as a ring of radius +1, where the radius is given in the ‘tubule radius’ (small feature) or ‘sheet radius’ (large feature) parameters. These convolution kernels were applied pixel-wise to determine if a pixel is brighter than the mean intensity of the surrounding ring of pixels. The strength of this filter was fine-tuned by adjusting the ‘tubule strength’ or ‘sheet strength’ parameters. Additionally, segmented pixels had to be brighter than the background levels defined above. This procedure generated two segmented images per channel, which were combined to generate the ‘total ER’ mask for that channel. To define which regions in each channel represented sheets or tubules a morphological opening was applied, the degree of which was controlled by the parameter ‘trimming factor’. Features that remained in the Sec63 segmentation mask after morphological opening were provisionally designated as sheets, the remainder of the ER mask was designated tubules. Finally, areas that were sheet-like in the Rtn1 segmentation mask and overlapped with the Sec63 mask were designated tubular clusters. Tubular clusters were subtracted from provisional sheets and added to the tubules to obtain the final designation of sheets and tubules. The median size measurements of each class were taken from the whole cell population. Values from independent experiments were averaged and standard errors of the mean were calculated.

### Quantitative real-time PCR

Quantitative real-time PCR was done exactly as described (Schmidt et al., 2019).

### Growth assays

For growth assays in liquid culture, cells were grown to saturation and diluted to OD_600_ = 0.05. For each strain, 500 *µ*l culture was transferred into 48-well plates in triplicate and absorbance at 600 nm was measured at room temperature in 5-minute intervals for 24 hours using a Tecan Infinite M1000 Pro plate reader. The area under the curve was calculated with the R package Growthcurver (Sprouffske and Wagner, 2016) and used as a measure for cell growth. For the growth assays shown in Figure 9C, cells were grown to mid log phase, diluted as above and absorbance was measured at 30°C using a Tecan Spark Cyto plate reader. This experimental regime ensured that *Δhac1* mutants grew as well as wild-type cells. Growth assays on solid medium were done as described (Schuck et al., 2009) using SCD plates with and without 0.2 µg/ml tunicamycin (Merck). Plates were imaged after 1.5 days of growth at 30°C.

### Lipidomics

For each sample, approximately 2 x 10^8^ cells (10 ODs) were harvested from cultures grown to mid log phase and snap frozen in liquid nitrogen. Cells were disrupted with glass beads as above in 50 mM HEPES pH 7.5 containing 0.5 mM EDTA. Aliquots were subjected to acidic Bligh and Dyer lipid extractions in the presence of internal lipid standards added from a master mix containing 40 pmol d7-PC mix (15:0/18:1-d7, Avanti Polar Lipids), 25 pmol PI (17:0/20:4, Avanti Polar Lipids), 25 pmol PE and 15 pmol PS (14:1/14:1, 20:1/20:1, 22:1/22:1, semi-synthesized as described in Özbalci et al., 2013), 20 pmol DAG (17:0/17:0, Larodan), 20 pmol TAG (D5-TAG-Mix, LM-6000 / D5-TAG 17:0,17:1,17:1, Avanti Polar Lipids), 20 pmol PA (PA 17:0/20:4, Avanti Polar Lipids), 5 pmol PG (14:1/14:1, 20:1/20:1, 22:1/22:1, semi-synthesized as described in Özbalci et al., 2013), 40 pmol ergosteryl ester (15:0 and 19:0, semi-synthesized as described in Gruber et al., 2018) and 20 pmol t-Cer (18:0, Avanti Polar Lipids). Lipids recovered in the organic extraction phase were evaporated by a gentle stream of nitrogen. Prior to measurements, lipid extracts were dissolved in 10 mM ammonium acetate in methanol and transferred into Eppendorf twin.tec 96-well plates. Mass spectrometric measurements were performed in positive ion mode on an AB SCIEX QTRAP 6500+ mass spectrometer equipped with chip-based (HD-D ESI Chip, Advion Biosciences) nano-electrospray infusion and ionization (Triversa Nanomate, Advion Biosciences) as described (Özbalci et al., 2013). The following precursor ion scanning (PREC) and neutral loss scanning (NL) modes were used for the measurement of the various lipid classes: +PREC 184 (PC), +PREC282 (t-Cer), +NL141 (PE), +NL185 (PS), +NL277 (PI), + NL189 (PG), +NL115 (PA), +PREC 77 (ergosterol), +PREC379 (ergosteryl ester). Ergosterol was quantified following derivatization to ergosterol acetate in the presence of 200 pmol of the internal standard (22E)-Stigmasta-5,7,22- trien-3-beta-ol (Aldrich, R202967) using 100 *µ*l acetic anhydride/chloroform (1:12 v/v) overnight under argon atmosphere (Ejsing et al., 2009). Mass spectrometry settings: Resolution: unit, low mass configuration; data accumulation: 400 MCA; curtain gas: 20; Interface heater temperature: 60; CAD: medium; DR: open; separation voltage: 3800; DMO: -3; compensation voltage: 10; DMS temperature: 60°C. Data evaluation was done using LipidView (Sciex) and a software developed in house, ShinyLipids.

### Pah1 dephosphorylation and phosphorylation assays

Microsomes were prepared from strains lacking *PEP4*, *PRB1*, *PAH1* and additional genes as indicated. Removal of Pep4 and Prb1 was critical to prevent protein degradation during in vitro assays. Removal of Pah1 ensured that all microsome donor strains had the same ER morphology and avoided contamination of microsomes with Pah1. Five-hundred ODs of cells were washed once with 100 mM Tris pH 9.4 containing 10 mM NaN_3_, incubated in 100 mM Tris pH 9.4 containing 10 mM NaN_3_ and 10 mM DTT at 30°C for 10 min, pelleted, resuspended to 20 OD/ml in spheroplast buffer (50 mM Tris pH 7.5, 1 M sorbitol) and treated with 0.2 U/OD zymolyase 100T (Biomol, Hamburg, Germany) at 30°C for 10 min. The resulting spheroplasts were washed three times with ice-cold spheroplast buffer and resuspended to 100 OD/ml in hypo-osmotic lysis buffer (50 mM HEPES pH 7.5, 2 mM EDTA, 200 mM sorbitol, 1 mM PMSF, protease inhibitors). Lysates were homogenized with 40 strokes of a Dounce homogenizer with a clearance of 0.01 - 0.06 mm (Kimble Chase) and cleared twice by centrifugation at 3,000 xg at 4°C for 5 min. Total membranes were pelleted by centrifugation at 16,000 xg at 4°C for 15 min, washed once with one volume of hypo-osmotic lysis buffer and resuspended in 200 *µ*l membrane buffer (20 mM HEPES pH 6.8, 250 mM sorbitol, 150 mM potassium acetate, 5 mM magnesium acetate). To enrich ER-derived microsomes, 200 *µ*l membranes were loaded onto a 3.8 ml two-step sucrose gradient (1.2 /1.5 M sucrose in 20 mM HEPES pH 7.4, 50 mM potassium acetate, 1 mM DTT, 2 mM EDTA) and centrifuged at 150,000 xg at 4°C for 1.5 h. Microsomes were collected from the 1.2/1.5 M sucrose interphase, diluted with five volumes membrane buffer, pelleted at 16,000 xg for 15 min, washed once with one volume membrane buffer and once with one volume reaction buffer (50 mM Tris pH 7.1, 150 mM NaCl, 100 mM sodium acetate, 100 mM MgCl_2_, 1 mM DTT, 0.1 mg/ml BSA and protease inhibitors). Finally, microsomes were resuspended in 50 *µ*l reaction buffer and dispersed by sonication.

To isolate Pah1-FLAG, 10 ODs of cells were lysed as for standard western blotting in 20 mM Tris pH 8 containing 150 mM KCl, 5 mM MgCl_2_, 1% Triton X-100, 1 mM PMSF, protease inhibitors and phosphatase inhibitors (PhosSTOP, Roche). Lysates were cleared by centrifugation at 16,000 xg at 4°C for 2 min and Pah1-FLAG was immunoprecipitated with anti-FLAG agarose beads (Sigma) at 4°C for 30 min. Beads were washed three times with the above lysis buffer and once with reaction buffer. Pah1-FLAG was eluted twice with 50 *µ*l 0.2 mg/ml FLAG peptide in reaction buffer and eluates were pooled. Fifty *µ*l microsomes from different strains were incubated with 50 *µ*l immunoisolated Pah1-FLAG from *Δnem1* or *Δice2* cells at 30°C for 30 min, proteins were resolved by Phos-tag PAGE and analyzed by western blotting.

### Co-immunoprecipitation

Cells were spheroplasted, lysed and homogenized as above. Lysates were cleared at 500 xg at 4°C for 2 min and incubated with 0.5% Triton X-100 at 4°C for 30 min. Insoluble material was removed at 100,000 xg for 30 min and Spo7-FLAG or Nem1- FLAG were precipitated from the supernatant with anti-FLAG agarose beads for 30 min. Beads were washed twice with hypo-osmotic lysis buffer containing 0.5% Triton X-100 and once with lysis buffer without detergent. Bound protein was eluted with SDS sample buffer at 65°C for 5 min and analyzed by western blotting.

### Proximity-dependent biotinylation assay

Cells were harvested and lysed as for standard western blotting but in SDS/Triton buffer (50 mM Tris pH 7.5, 0.4% SDS, 2% Triton X-100, 150 mM NaCl, 5 mM EDTA, 1 mM DTT, protease inhibitors). Biotinylated proteins were captured with streptavidin agarose beads (Thermo Fisher) at 4°C for 1 h. Beads were washed twice with SDS/Triton buffer at 4°C for 5 min, twice with 10% SDS at room temperature for 5 min, twice with 50 mM HEPES pH 7.4 containing 500 mM NaCl, 1 mM EDTA, 1% Triton X- 100 and 0.1% sodium deoxycholate, and twice with 50 mM Tris pH 7.5 containing 50 mM NaCl and 0.1% Triton X-100. Bound protein was eluted with SDS-PAGE sample buffer containing 2 mM biotin at 65°C for 10 min and analyzed by western blotting.

## ACKNOWLEDGEMENTS

We thank Randy Schekman and Symeon Siniossoglou for reagents; the flow cytometry & FACS and imaging facilities at the ZMBH for assistance; Oliver Pajonk, Dorottya Polos and Julia Schessner for early contributions to this project; Iris Leibrecht for help with lipidomics analyses; Marie-Pierre Péli-Gulli and Emmanuelle Dubots for advice on Phos-tag gels; Rose Goodchild, Daniel Markgraf, Nicolai Karcher, Robin Klemm, Savvas Paragkamian, Sophie Winter and all Schookees for comments on the manuscript. This work was supported by a fellowship from the Heidelberg Biosciences International Graduate School to DP and grant EXC81 from the Deutsche Forschungsgemeinschaft (DFG). BB was funded by projects 319506281-TRR 186 and 112927078-TRR 83 from the DFG. The authors declare no competing financial interests.

## AUTHOR CONTRIBUTIONS

Conceptualization: Peter Bircham, Dimitrios Papagiannidis, Sebastian Schuck; Formal analysis: Peter Bircham; Investigation: Peter Bircham, Christian Lüchtenborg, Dimitrios Papagiannidis, Giulia Ruffini; Software: Peter Bircham; Supervision: Britta Brügger, Sebastian Schuck; Writing - original draft: Dimitrios Papagiannidis, Sebastian Schuck; Writing - review and editing: all authors.

---

**Figure S1.**
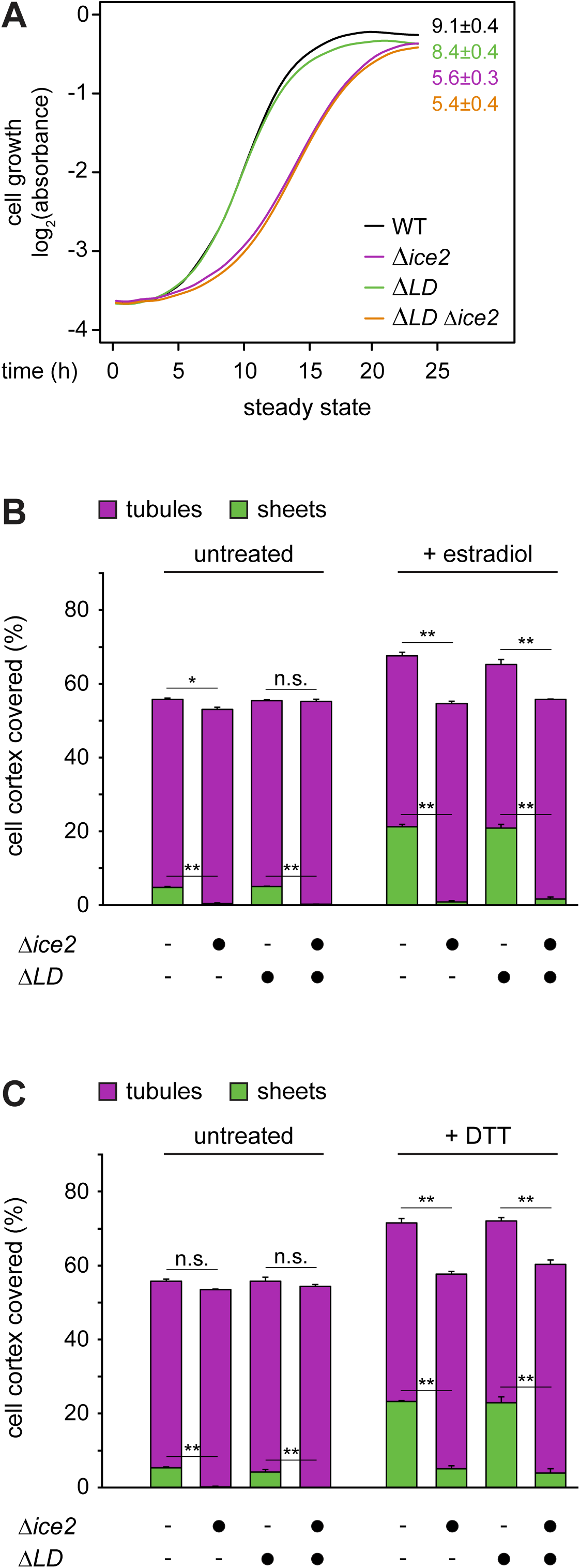
Absence of lipid droplets has no effect on ER expansion in WT or *Δice2* cells. **(A)** Growth assays of untreated WT, *Δice2*, *ΔLD* and *ΔLD Δice2* cells (SSY2228, 2229, 2230, 2256). Numbers represent areas under the curves and serve as growth indices. Mean ± SEM, n = 3. *ΔLD*, *Δlipid droplet*. **(B)** Quantification of peripheral ER structures in WT, *Δice2*, *ΔLD* and *ΔLD Δice2* cells harboring the inducible system (SSY2598, 2599, 2600, 2601), which were untreated or treated with 800 nM estradiol for 6 h. Plotted is the mean percentage of cell cortex covered by tubules (purple) or sheets (green), n = 3. Upper error bars are SEM for the sum of tubules and sheets, lower error bars are SEM for sheets. Asterisks indicate statistical significance. *, p < 0.05; **, p < 0.01; n.s., not significant. **(C)** As in panel B but after treatment with 8 mM DTT for 1 h.

**Table S3.** Plasmids used in this study. GEM = GAL4DBD-EstR-Msn2TAD.

| Plasmid | Alias | Source |
| --- | --- | --- |
| pRS415-P<sub>MET25</sub>-ino2(L119A) | pSS108 | Heyken et al., 2005 |
| pRS416-P<sub>GAL1</sub> | pSS031 | Mumberg et al., 1994 |
| pRS415-P<sub>GAL1</sub>-ino2(L119A) | pSS448 | this study |
| pNH605-P<sub>ADH1</sub>-GEM | pDEP151 | David Pincus |
| pNH605-P<sub>ADH1</sub>-GEM-P<sub>GAL1</sub> | pSS474 | Schmidt et al., 2019 |
| pNH605-P<sub>ADH1</sub>-GEM-P<sub>GAL1</sub>-ino2(L119A) | pSS475 | this study |
| pRS306-P<sub>ADH1</sub>-GEM | pDEP001 | Pincus et al., 2014 |
| pRS306-P<sub>ADH1</sub>-GEM-P<sub>GAL1</sub> | pSS476 | Schmidt et al., 2019 |
| pRS306-P<sub>ADH1</sub>-GEM-P<sub>GAL1</sub>-ino2(L119A) | pSS477 | this study |
| pRS303H-P<sub>GPD</sub>-TagBFP | pMaM245 | Szoradi et al., 2018 |
| pFA6a-mNeon-kanMX4 | pMaM375 | Michael Knop |
| pFA6a-GFP(S65T)-kanMX6 | pSS037 | Longtine et al., 1998 |
| pFA6a-GFP(S65T)-HISMX6 | pSS039 | Longtine et al., 1998 |
| pFA6a-mNeon-kanMX6 | pSS445 | Schäfer et al., 2020 |
| pFA6a-mNeon-HIS3MX6 | pSS447 | this study |
| pRS415-P<sub>ADH1</sub> | pSS022 | Mumberg et al., 1995 |
| pRS415-P<sub>ADH1</sub>-Ice2 | pSS761 | this study |
| pRS304-4xUPRE-GFP | pDEP017 | Pincus et al., 2010 |
| pFA6a-3xHA-HISMX6 | pSS042 | Longtine et al., 1998 |
| pFA6a-3xFLAG-kanMX6 | pSS072 | this study |
| pFA6a-TurboID-3myc-kanMX6 | pSS1061 | Larochelle et al., 2019 |
| YCplac111-Pah1-PrtA | pSS1005 | O'Hara et al., 2006 |
| YCplac111-pah1(7A)-PrtA | pSS1006 | O'Hara et al., 2006 |
| YCplac111-Pah1-3HA | pSS1045 | this study |
| YCplac111-pah1(7A)-3HA | pSS1047 | this study |

**Table S4.**
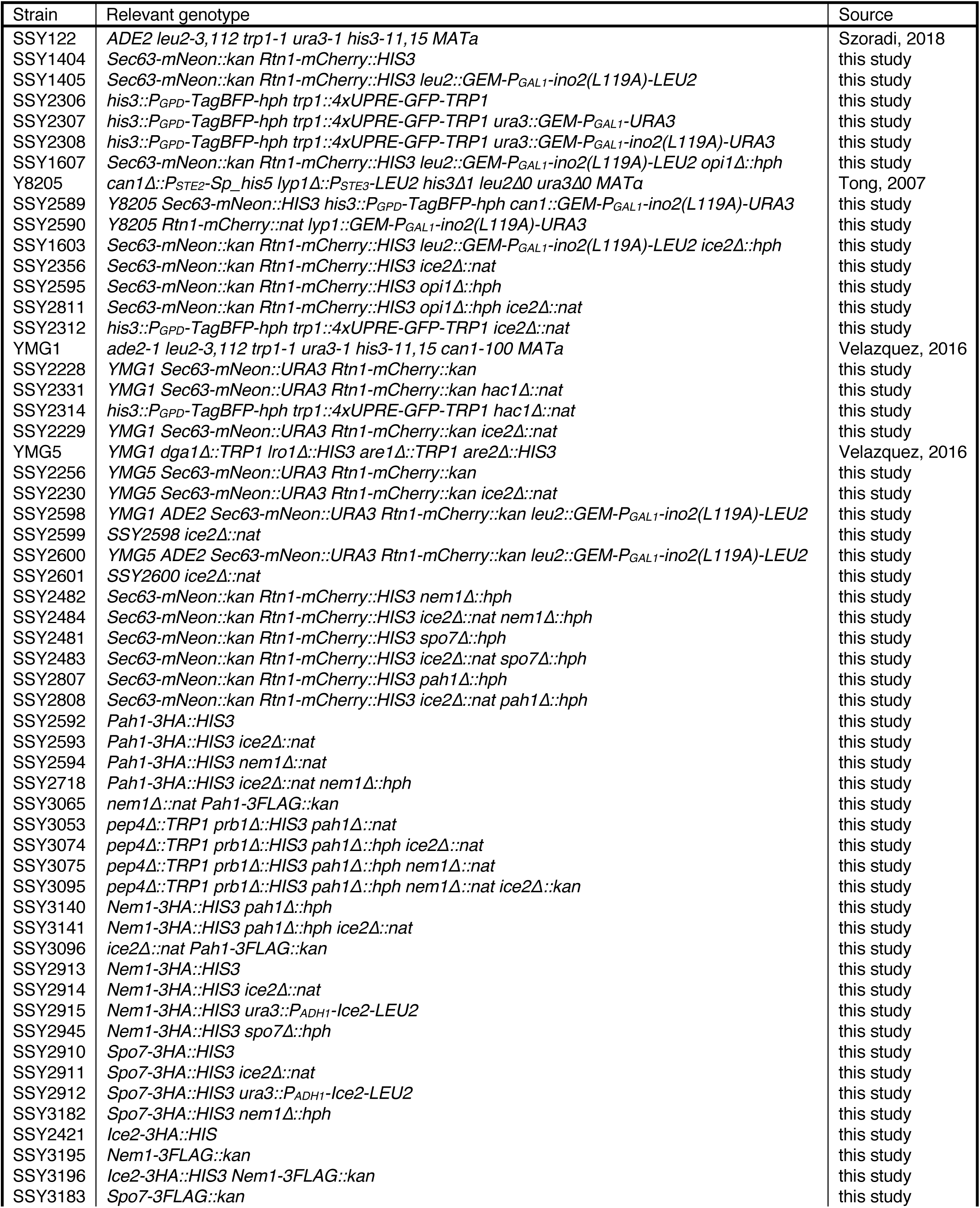
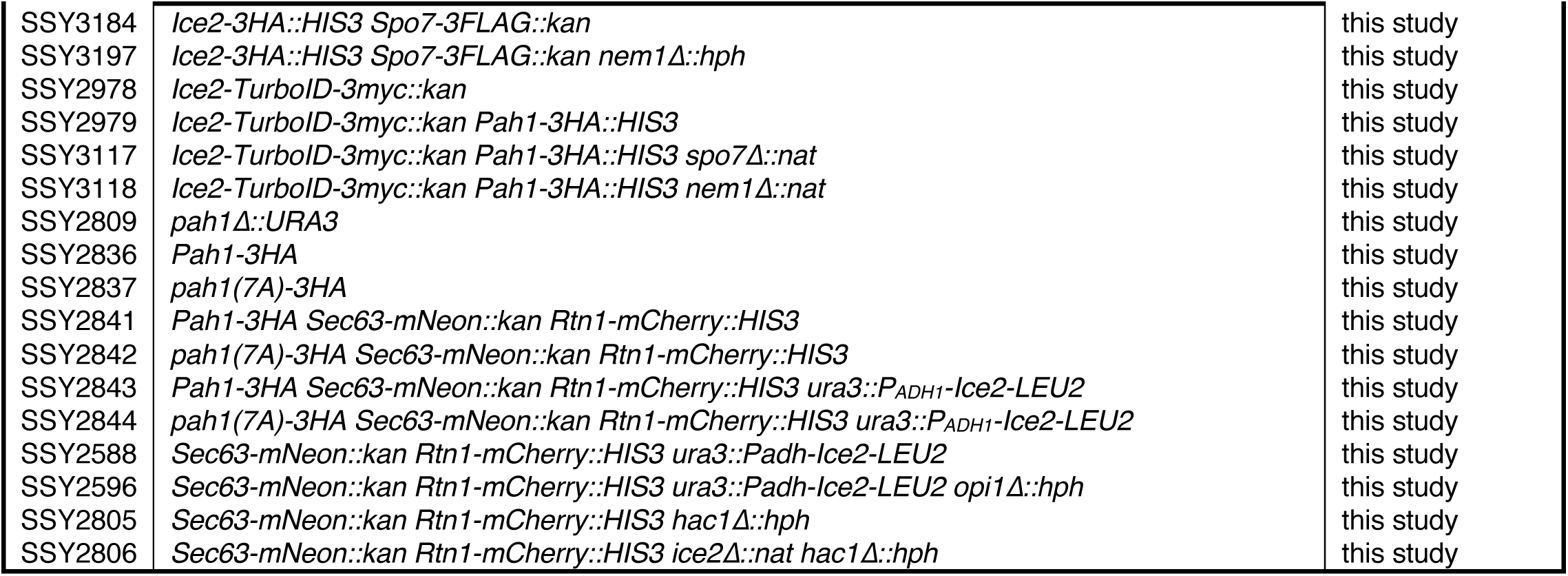
Yeast strains used in this study. mNeon = mNeonGreen. GEM = P_ADH1_- GAL4DBD-EstR-Msn2TAD.

## Notes

### Competing Interest Statement

The authors have declared no competing interest.

